# Evolutionary co-option of the ancient protein AMBRA1 in plant photomorphogenesis

**DOI:** 10.64898/2026.10.14.758420

**Authors:** Claudia Beraldo, Vasilii Apollonov, Luisa Dalla Valle, Luis Vidali, Andreas Hiltbrunner, Tomas Morosinotto, Alessandro Alboresi

**Affiliations:** Dipartimento di Biologia, Università di Padova, Via Ugo Bassi 58/B, 35121, Padova; Department of Biology and Biotechnology and Bioinformatics and Computational Biology Program, Worcester Polytechnic Institute, Worcester, Massachusetts, United States of America; Faculty of Biology, University of Freiburg, 79104, Freiburg, Germany; CIBSS - Center for Integrative Biological Signalling Studies, University of Freiburg, 79104, Freiburg, Germany

**Keywords:** light signalling, AMBRA1, photomorphogenesis, evolution

## Abstract

Photomorphogenesis, the light-driven developmental program of plants, is repressed in darkness by the CUL4-DDB1-COP1 ubiquitin ligase complex. In animals, AMBRA1 is a substrate receptor for the same CUL4-DDB1 machinery, regulating proteostasis, cell proliferation and autophagy, but its role in plants has remained unknown. Given that COP1 and AMBRA1 are both substrate receptors of CUL4–DDB1, we asked whether AMBRA1 contributes to the repression of photomorphogenesis in plants.

Here we show that the moss *Physcomitrium patens* carries two light-regulated AMBRA1 genes, *AMBRA1a* and *AMBRA1b*, whose loss impairs vegetative growth and cell-fate transitions under light and prevents the elongation of gravitropic filaments in darkness, indicating that AMBRA1 is required to repress photomorphogenesis. AMBRA1 localizes to the nucleus under both long-day and dark conditions, and its loss causes widespread transcriptional reprogramming of developmental and metabolic pathways, including repression of photosynthesis-related genes in darkness. These findings identify AMBRA1 as a previously unrecognized nuclear regulator that links light availability to plant growth and as a key repressor of photomorphogenesis.

## Introduction

Land plants are sessile organisms that cannot escape unfavourable conditions and must adjust their growth to environmental changes^1,2^. Light is a particularly critical cue, acting both as a developmental signal and as the energy source for photosynthesis. Plants have evolved dedicated photoreceptors and signalling pathways that couple light perception to developmental programmes. Light promotes photomorphogenesis, the developmental programme that builds photosynthetically competent tissues, while darkness suppresses it in favour of skotomorphogenic growth^3^. This developmental switch depends on tightly regulated proteostasis. In darkness, the E3 ubiquitin ligase CONSTITUTIVE PHOTOMORPHOGENIC 1 (COP1) associates with SUPPRESSOR OF PHYA-105 (SPA) proteins and the CULLIN4 (CUL4)-DDB1 machinery to promote the degradation of positive regulators of light signaling, including ELONGATED HYPOCOTYL 5 (HY5) and several photoreceptors^4–6^. CUL4-DDB1 is an ancient scaffold conserved across kingdoms: in animals, related CUL4-DDB1 complexes recruit distinct substrate receptors to control processes ranging from DNA repair and cell-cycle progression to proteostasis^7–9^. COP1 also has an animal counterpart and, in mammals, participates with DE-ETIOLATED 1 (DET1), another protein conserved between plants and animals, in a CUL4-DDB1-containing ubiquitin-ligase complex^10,11^. This conservation raises the possibility that plants and animals share additional substrate receptors that recruit specific proteins to CUL4-DDB1 complexes for ubiquitination and degradation.

Activating Molecule in Beclin1-Regulated Autophagy 1 (AMBRA1) is a candidate for such a conserved regulator because it is widely conserved across eukaryotes and regulates ubiquitin-dependent protein turnover in animals. AMBRA1 is a large, intrinsically disordered protein in which conserved N- and C-terminal regions fold into a WD40 domain that mediates its interaction with DDB1 and provides a site for substrate recruitment^12–15^. Within the CUL4-DDB1 ubiquitin-ligase complex, in animals AMBRA1 acts as a substrate receptor, directly recruiting proteins such as cyclin D2 for ubiquitylation and proteasomal degradation, thereby restricting cell-cycle progression^14–16^. AMBRA1 is additionally implicated in autophagy initiation via the Beclin1-PI3KC3 complex^17–20^.

Despite the conservation of AMBRA1 homologues and their characteristic WD40-containing architecture in plant genomes, their biological functions in plants remain unknown. Here, we combine comparative genomics, cell biology, genetics and transcriptomics to investigate the functions of the two AMBRA1 homologues, AMBRA1a and AMBRA1b, in the model moss *Physcomitrium patens*. We show that they are nuclear proteins required for protonemal cell-fate transitions, vegetative growth and gravitropic responses, as well as for the repression of photomorphogenesis in darkness. Loss of AMBRA1 results in widespread transcriptional reprogramming, affecting developmental, cell-wall, cell-cycle and photosynthesis-associated pathways. Comparative phenotypic and transcriptomic analyses with *cop1* null mutants revealed only partial convergence in the repression of photomorphogenesis and in the regulation of photosynthesis-associated genes in darkness. Together, our findings identify AMBRA1 as a previously uncharacterized nuclear protein required for plant development and the repression of photomorphogenesis in darkness.

## Results

### Plant AMBRA1 proteins show a conserved domain architecture and light-responsive expression

To determine the evolutionary distribution of AMBRA1, we searched animal, plant and algal genomes using human AMBRA1 (HsAMBRA1) as the query. AMBRA1 homologues were identified outside animals in red algae, stramenopiles, green algae, bryophytes and vascular plants (Supplementary Table 1). Maximum-likelihood phylogenetic analysis, using the related WD40 proteins KATNB1 and WDR5 as outgroups, separated the AMBRA1 homologues of Viridiplantae, Rhodophyta and Stramenopiles from metazoan AMBRA1 (Fig. 1a). The position of algal sequences was consistent with the presence of AMBRA1 before the emergence of land plants. AMBRA1 copy number was low but varied among lineages: several algal and land-plant species contained a single homologue, whereas *Physcomitrium patens* and most sampled angiosperms contained two or three copies. Some duplications were lineage-specific, such as the three AMBRA1 homologues of the hexaploid *Triticum aestivum*, which group together (Fig. 1a). In other cases, such as *Zea mays* and *Sorghum bicolor* or *Lotus japonicus* and *Medicago truncatula*, AMBRA1a and AMBRA1b copies group by paralogue rather than by species (Fig. 1a).

**Figure 1.**
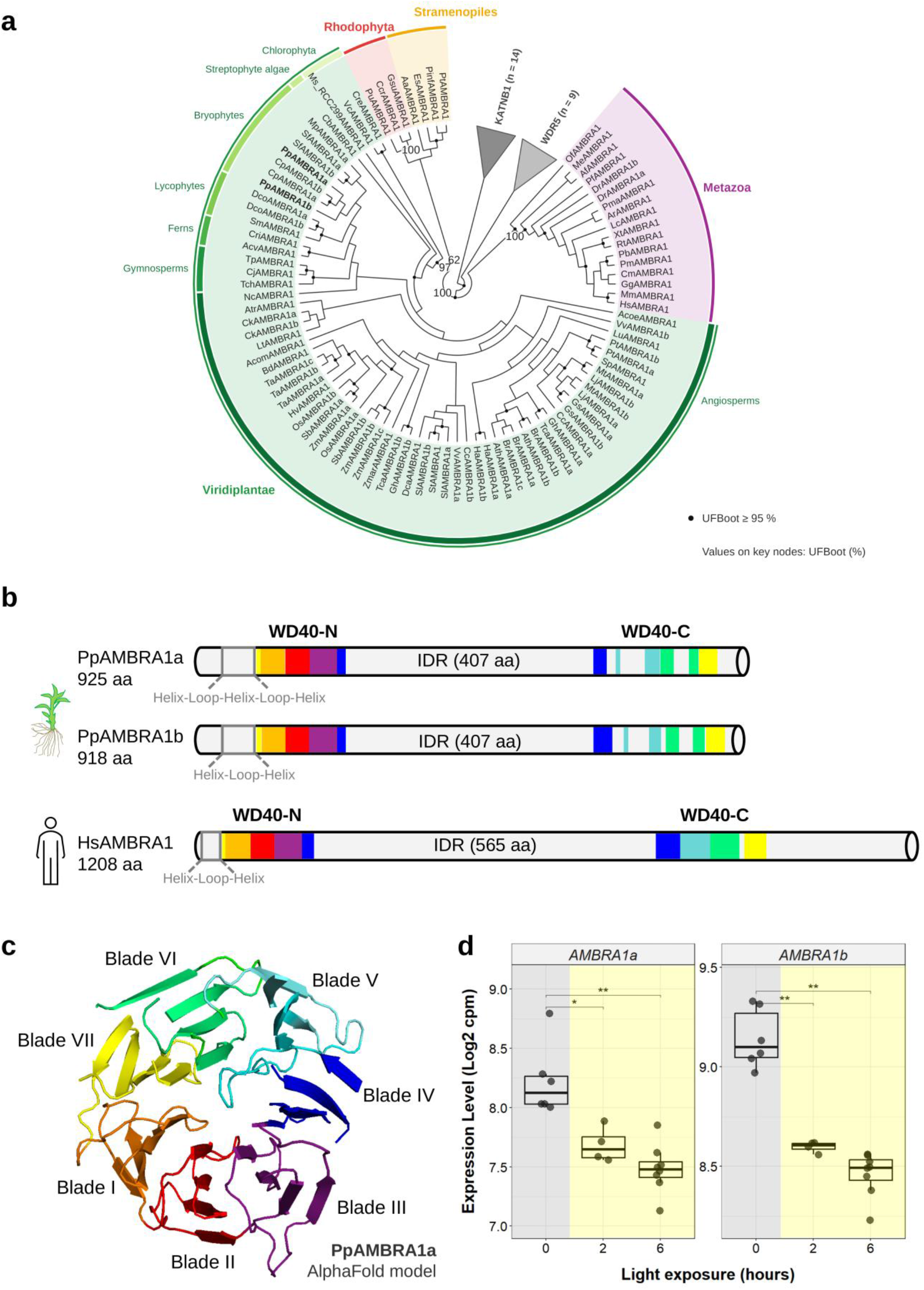
Evolutionary and structural analysis of AMBRA1 proteins and time-resolved expression of *AMBRA1* genes in *Physcomitrium patens*. a) Maximum-likelihood phylogeny of AMBRA1 proteins from Metazoa, Stramenopiles, Rhodophyta and Viridiplantae (Chlorophyta, streptophyte algae, bryophytes, lycophytes, ferns, gymnosperms and angiosperms), shown as a cladogram (branch lengths not to scale). KATNB1 and WDR5 sequences (collapsed) were used as outgroups and the tree was rooted on the KATNB1 clade. Sequences are listed in Supplementary Table 1. Black dots indicate nodes with ultrafast bootstrap (UFBoot) support ≥ 95%; numbers at key nodes indicate UFBoot values (%). *P. patens* AMBRA1a and AMBRA1b are shown in bold. b) Schematic representation of the *P. patens* AMBRA1a, AMBRA1b and *H. sapiens* AMBRA1 proteins. The seven blades of the WD40 β-propeller structure, each composed of four antiparallel β-strands, are highlighted in different colours. c) Alphafold model prediction of *P. patens* AMBRA1a WD40 core domain (AlphaFold ID: A0A2K1JYD0-F1). d) Gene expression regulation of *P. patens AMBRA1a* and *AMBRA1b* under light exposure. Boxplots show expression levels Log_2_cpm at 0, 2, and 6 hours of light treatment. Grey and yellow backgrounds indicate darkness and light, respectively. Individual data points represent biological replicates from a previous transcriptomic experiment^21^. Statistical significance between time points was evaluated using Wilcoxon rank-sum tests; only significant comparisons are displayed with brackets (* p < 0.05, ** p < 0.01).

We next examined whether the two *P. patens* homologues, hereafter AMBRA1a and AMBRA1b, retained the structural features of human AMBRA1. AlphaFold models predicted that AMBRA1a and AMBRA1b contained a seven-bladed β-propeller in which conserved N- and C-terminal regions contributed to a discontinuous WD40 domain separated by an extensive intrinsically disordered region, as described for HsAMBRA1 (Fig. 1b-c, Supplementary Fig. 1 and 2). The highest sequence conservation was detected within the N-terminal portion of the WD40 domain (Supplementary Fig. 1). In HsAMBRA1, an N-terminal helix-loop-helix motif contributes to DDB1 binding^16^. The corresponding regions of AMBRA1a and AMBRA1b were predicted to retain a related helical organization but contained expanded loop regions (Supplementary Fig. 3). Structural superposition nevertheless indicated conservation of the overall N-terminal fold (RMSD in all comparisons <1Å) among HsAMBRA1, AMBRA1a and AMBRA1b.

**Figure 2.**
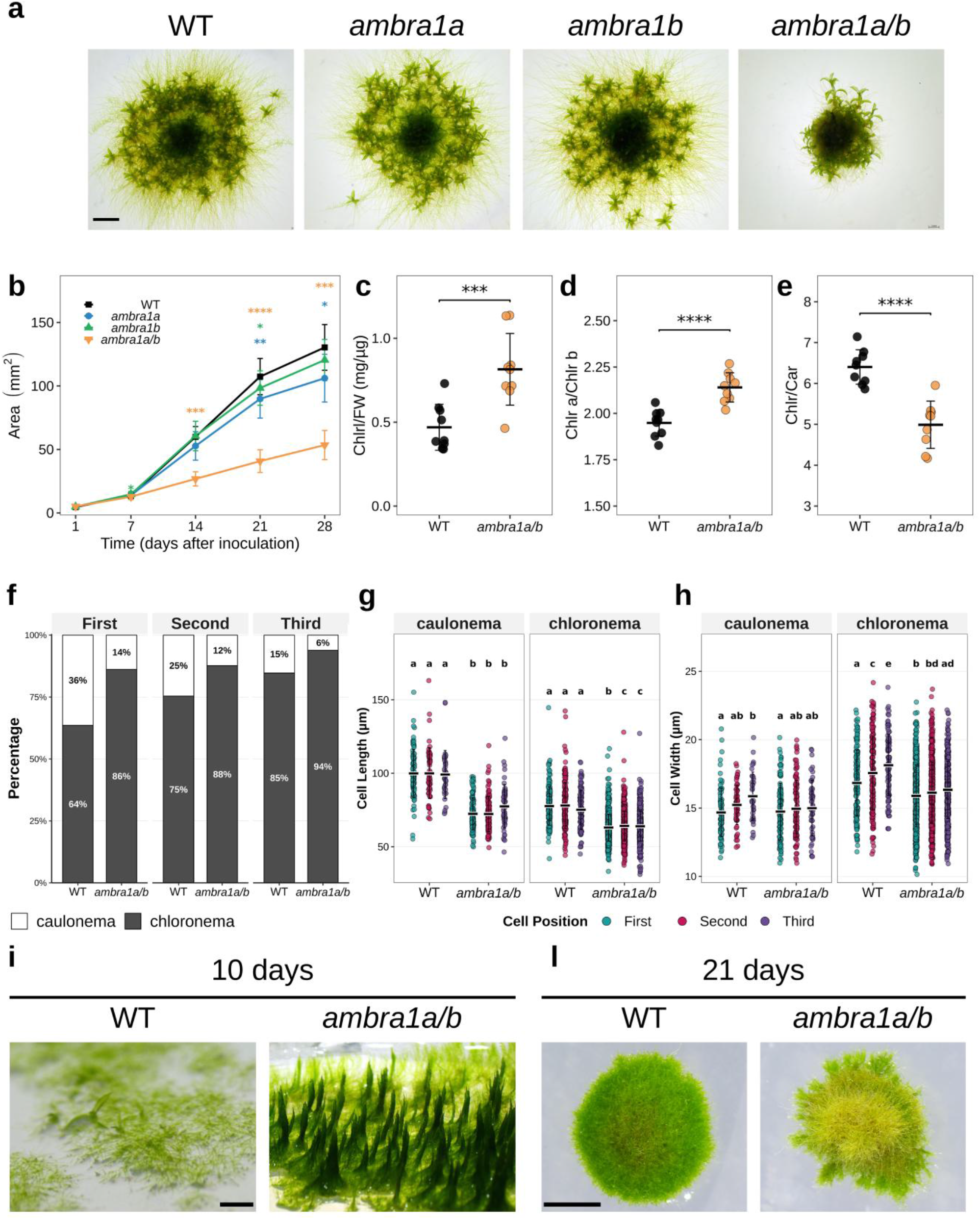
Role of AMBRA1 in *Physcomitrium patens* plant growth under long day conditions. a) Representative images of *P. patens* colonies grown horizontally on minimal medium for 28 days under standard photoperiod conditions (LD). Scale bar = 2 mm. b) Colony area was measured weekly over a 28-day period. n = 3 independent experiments, each with 8-24 colonies per genotype. Asterisks are color-coded by genotype (blue: *ambra1a*; green: *ambra1b*; orange: *ambra1a/b*) and indicate statistically significant differences compared to WT (Student’s t-test or Wilcoxon rank-sum test depending on data distribution, assessed by Shapiro-Wilk test; *p<0.05, **p<0.01, ***p<0.001, ****p<0.0001). c-e) Quantification of total chlorophyll (Chlr) content per fresh weight (FW) (c), Chlr *a/* Chlr *b* ratio (d) and Chlr / carotenoid (Car) (e) ratio in WT and *ambra1a/b* double-mutant lines. Each dot represents a biological replicate. Asterisks indicate statistically significant differences compared to WT (two-tailed Student’s t-test or Wilcoxon rank-sum test, depending on data normality as assessed by Shapiro-Wilk test; ** p<0.01, *** p<0.001, **** p<0.0001). f) Chloronema-to-caulonema ratio (percentage from the first, second, and third cell of each filament) in WT and *ambra1a/b* KO. g-h) Cell length (g) and cell width (h) of caulonema and chloronema filaments in WT and *ambra1a/b* mutants from the first, second, and third cell of each filament. Groups not sharing a letter are significantly different (Kruskal-Wallis test with Dunn’s post-hoc test, BH-adjusted, p<0.05). i-l) Pictures of *P. patens* colonies grown on medium supplemented with glucose and ammonium tartrate for 10 days (i) or 21 days (l) under standard photoperiod conditions. Scale bar = 2 mm.

We then examined the expression of *AMBRA1a* and *AMBRA1b* during the transition from darkness to light. Transcripts of both genes accumulated in darkness and declined rapidly following illumination (Fig. 1d). A similar expression pattern was observed for the two *Arabidopsis thaliana* homologues in published datasets (Supplementary Fig. 4). Co-expression analysis using a set of 38 photomorphogenesis-related genes^9^ further showed that *AMBRA1a* and *AMBRA1b* expression covaried with several components of light signalling (Supplementary Fig. 5, Supplementary Table 2). Both genes were positively correlated with *PpCUL4a* and *PpCUL4c*, while *PpHY5b* and *PpPIF3* were among the strongest positive correlations. Thus, the conserved architecture and light-responsive expression of plant AMBRA1 homologues provided the rationale for testing their functions in light-dependent development.

### AMBRA1 is required for colony growth and protonemal cell-fate transitions under long-day conditions

To investigate the functional role of AMBRA1 in *P. patens*, we generated independent single KO lines (*ambra1a* and *ambra1b*) and a double KO (*ambra1a/b*) via gene targeting mediated by homologous recombination. The targeted loci and loss of the corresponding transcripts were verified, and independent lines of each genotype showed consistent phenotypes (Supplementary Fig. 6).

Under long day conditions and minimum medium, the *ambra1a/b* double mutant displayed more compact colony morphology (Fig. 2a) quantified as a reduction in colony area (Fig. 2b) and fresh weight (Supplementary Fig. 7) compared to both the WT and single mutants. Single *ambra1a* and *ambra1b* mutants showed milder reductions in colony area at later timepoints, indicating partial functional redundancy between the two paralogs, which was more fully revealed in the double mutant. In addition, the double mutant accumulated significantly higher chlorophyll levels than the WT and showed increased chlorophyll *a*/*b* ratio and a decreased chlorophyll-to-carotenoid ratio (Fig. 2c-e).

To identify the cellular basis of the growth defect, we examined protonemal development. The protonema is a tissue of mosses, consisting of a network of filaments that allows colony expansion. It comprises chloronemal cells, which contain numerous chloroplasts and are primarily photosynthetic, and faster-growing caulonemal cells, which promote radial colony expansion^22^. The *ambra1a/b* mutant had a higher chloronema-to-caulonema ratio than WT across the first three subapical cell positions, indicating an impaired transition between these two cell types (Fig. 2f). Compared with WT, both chloronemal and caulonemal cells were shorter in the mutant (Fig. 2g), and chloronemal cells also showed a reduction in width (Fig. 2h). The reduced cell dimensions provide a cellular explanation for the impaired growth of *ambra1a/b* colonies, whereas the increased proportion of chloroplast-rich chloronemal cells explains their higher chlorophyll content.

To test whether additional nutrient supply could rescue the phenotype, we grew *ambra1a/b* and WT plants on medium supplemented with glucose and ammonium tartrate. At early stages of growth, *ambra1a/b* protonemata adopted a pronounced vertical orientation, whereas WT protonemata grew along the surface of the medium (Fig. 2i). After 21 days, *ambra1a/b* colonies remained smaller than WT and developed a paler, senescence-like appearance (Fig. 2l).

Overall, these results establish that AMBRA1 is required for colony growth and protonemal cell-fate transitions under long-day conditions.

### AMBRA1 is required for the repression of photomorphogenesis in darkness

When plants were grown on vertical plates in continuous darkness, WT and single-mutant lines developed long, gravitropic filaments called skotonemata. *ambra1a* and *ambra1b* showed mild reductions in skotonemata density and length relative to WT, while the *ambra1a/b* double mutant produced few skotonemata, which failed to elongate (Fig. 3a-c). High-magnification imaging of individual filament tips further revealed increased branching in dark-grown *ambra1a/b* filaments (Fig. 3d). Mutant filaments also appeared greener than WT both in darkness and after 24 h of light exposure (Fig. 3d). Consistent with this observation, pigment quantification showed that *ambra1a/b* accumulated more chlorophyll per unit fresh weight than WT even in complete darkness and exhibited a decreased chlorophyll-to-carotenoid ratio (Fig. 3e-g).

**Figure 3.**
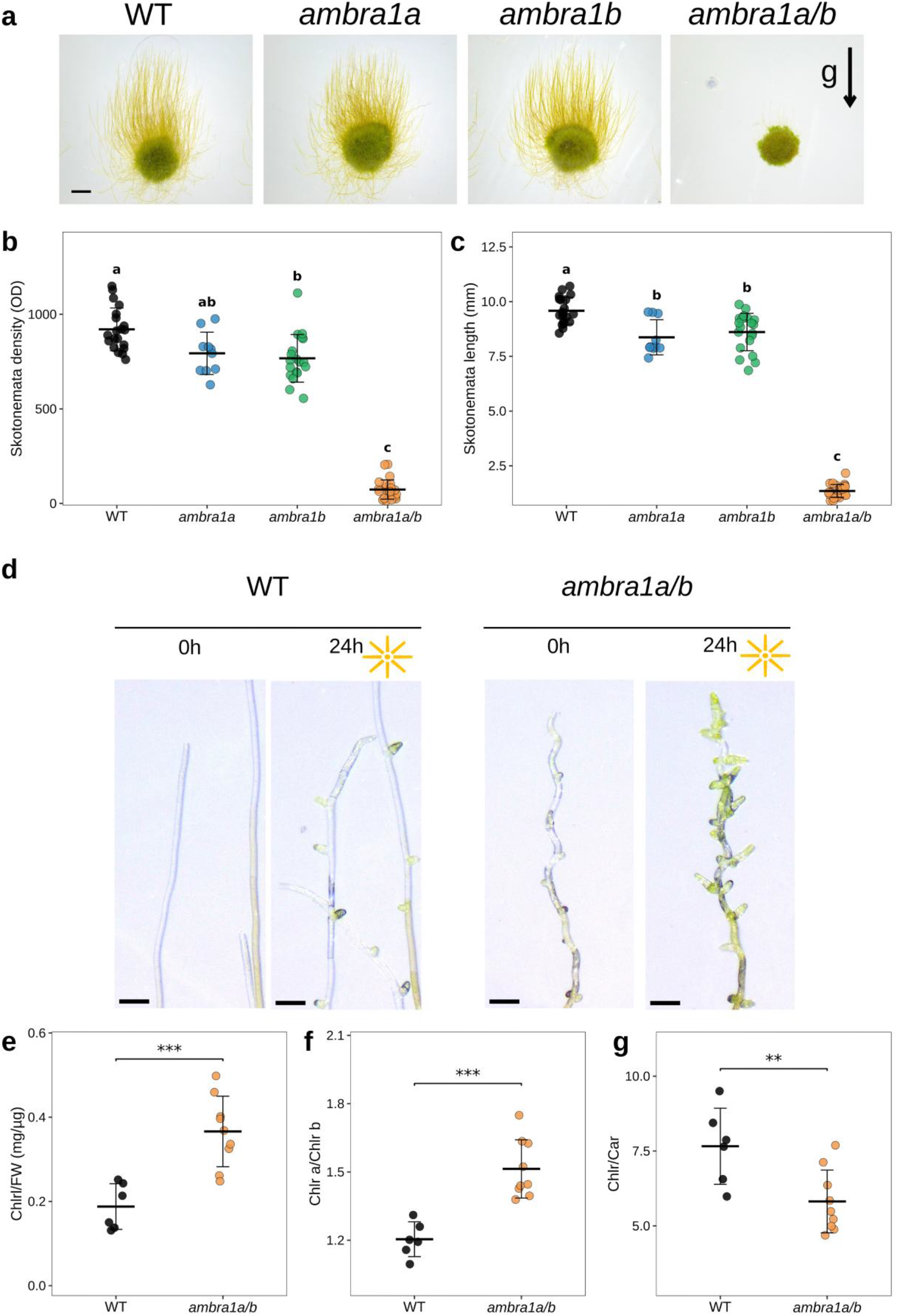
Role of AMBRA1 in *Physcomitrium patens* plant growth under continuous darkness. a) Representative images of *P. patens* colonies grown vertically on rich medium in continuous darkness for 14 days to trigger dark-induced development. Scale bar = 2 mm. The arrow (*g*) indicates the direction of the gravity vector. b) Skotonemata density of colonies grown in continuous darkness for 14 days, calculated as the optical density (OD) of the region above the colony edge, with width equal to the colony diameter and height of 1 mm. Each dot represents one colony (n ≥10). Statistical significance was assessed using a Kruskal-Wallis test followed by Dunn’s post-hoc test. Groups sharing the same letter are not significantly different (p ≥ 0.05). c) Skotonemata length (mm) of colonies grown in continuous darkness for 14 days, calculated as the mean of 6 skotonemata per colony (n ≥10). Statistics as in b). d) Representative images of individual filament tips from WT and *ambra1a/b* colonies grown in continuous darkness, imaged before (0h) and after 24h of light exposure Scale bar = 100µm. e-g) Quantification of total chlorophyll content per fresh weight (Chlr/FW; e), chlorophyll *a*/*b* ratio (f) and chlorophyll-to-carotenoid ratio (g) in WT and *ambra1a/b* double-mutant lines grown in continuous darkness. Each dot represents a biological replicate. Asterisks indicate statistically significant differences compared to WT (two-tailed Student’s t-test or Mann-Whitney/Wilcoxon rank-sum test for non-normally distributed data, assessed by Shapiro-Wilk test; ** p<0.01, *** p<0.001, **** p<0.0001).

Marker-free *ambra1a/b* lines, obtained by Cre-lox excision of the resistance cassettes, showed the same developmental and pigment defects as the parental lines, confirming that these phenotypes were not due to the resistance cassettes (Supplementary Fig. 8).

The failure to elongate skotonemata and the retention of chlorophyll in darkness show that AMBRA1 is required for the repression of photomorphogenic traits in the absence of light.

### AMBRA1 localizes to the nucleus

To gain insight into the functional role and subcellular localization of AMBRA1, we amplified *AMBRA1a* and *AMBRA1b* coding sequences from moss cDNA and then expressed full-length AMBRA1a and AMBRA1b proteins with a C-terminal Yellow Fluorescent Protein (YFP) tag, under the control of the constitutive PpEF1α promoter in the *ambra1a/b* double mutant background. As a prerequisite for interpreting their localization, we first verified that both fusion proteins were functional by testing their ability to complement the growth defects of *ambra1a/b*. The expression of AMBRA1b-YFP successfully restored filament elongation to WT levels (Fig. 4a), while the AMBRA1a-YFP fusion partially rescued the *ambra1a/b* mutant phenotype in darkness.

**Figure 4.**
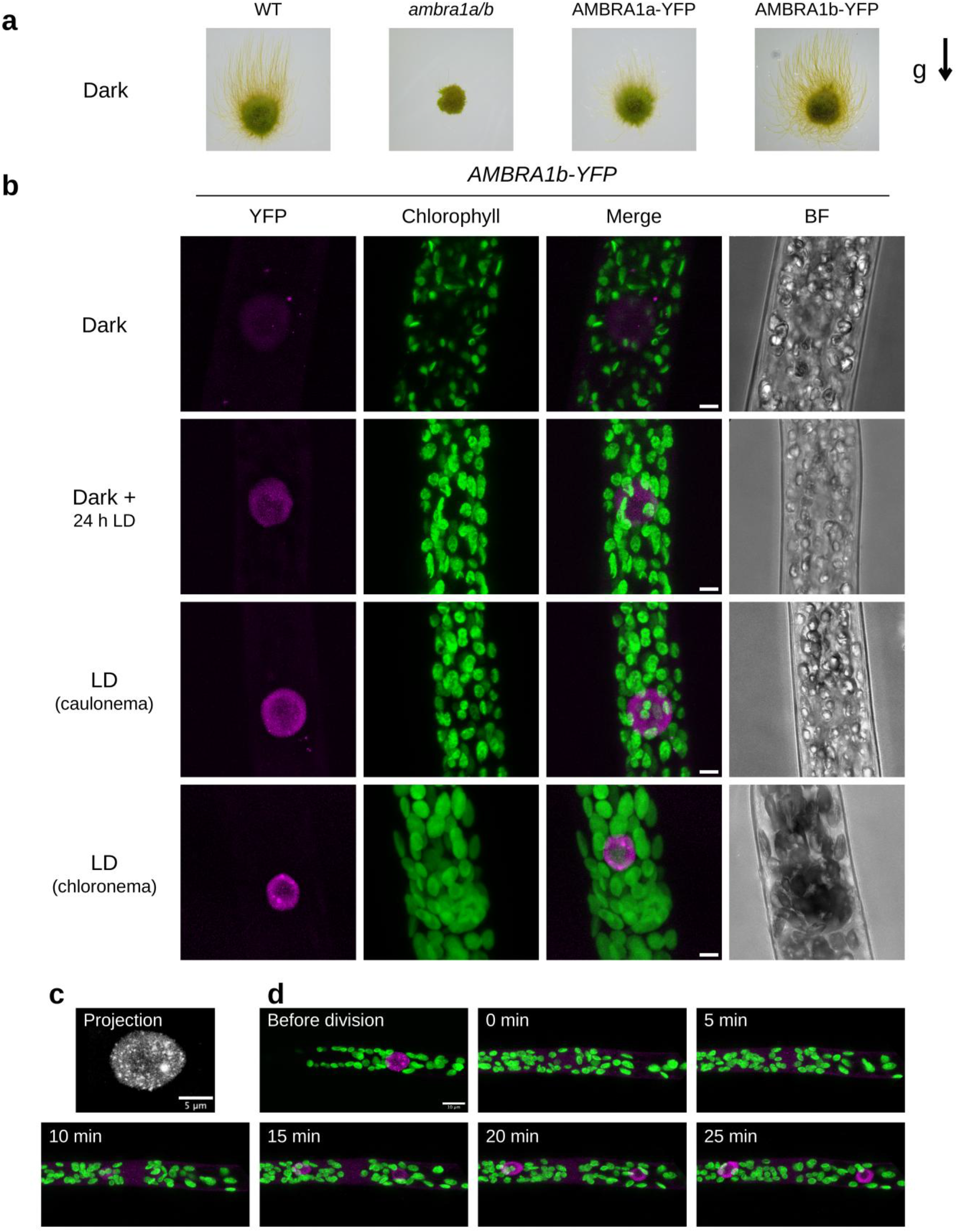
AMBRA1 localizes to the nucleus. a) Representative images of WT, *ambra1a/b* double knockout, and *ambra1a/b* lines transformed with AMBRA1a-YFP or AMBRA1b-YFP constructs driven by the PpEF1α promoter (PpEF1α::AMBRA1a-YFP and PpEF1α::AMBRA1b-YFP), grown vertically in the dark for 14 days. Scale bar = 5 mm. The arrow (g) indicates the direction of the gravity vector. b) Subcellular localization of AMBRA1b-YFP in protonemal cells grown in continuous darkness (Dark), after 24 h of light following darkness (Dark + 24 h LD), and in caulonemal and chloronemal cells grown under long-day conditions (LD). Green represents chlorophyll autofluorescence, magenta corresponds to YFP signal and merged, brightfield images are shown on the right. Scale bars = 5 µm. c) Maximum-intensity projection of a high-magnification confocal image stack of a nucleus showing the punctate/foci-like subnuclear distribution of AMBRA1b-YFP. Scale bar = 5 µm. d) Time-lapse frames of nuclear division showing AMBRA1 localization before division and during mitosis, imaged over a 25-minute period (0, 5, 10, 15, 20, 25 min; see also Supplementary Movie 1). Scale bars = 10 µm. The nucleus in C is from the same cell as in D.

Confocal laser-scanning microscopy showed that AMBRA1b-YFP localized predominantly to the nucleus in protonemal cells across all conditions examined, such as continuous darkness, 24 h after transfer to light, and long-day-grown caulonemal and chloronemal cells (Fig. 4b), as well as in bud and phyllid cells (Supplementary Fig. 9). Consistent with the more complete phenotypic complementation of AMBRA1b-YFP as compared to AMBRA1a-YFP, they also showed a brighter fluorescent signal (Fig. 4b and Supplementary Fig. 10). In general, the observed localization agreed with DeepLoc 2.0 predictions, which assigned AMBRA1a and AMBRA1b to the nucleus with probabilities of 0.7061 and 0.7230, respectively (Supplementary Table 3). High-resolution imaging further revealed that AMBRA1b-YFP accumulated in discrete regions within the nucleoplasm (Fig. 4b, c). These structures were more prominent in light-grown samples but remained detectable under continuous darkness (Fig. 4b). During cell division, the nuclear YFP signal was transiently lost and became dispersed throughout the cell, with the protein re-entering the newly formed daughter nuclei once division is complete. (Fig. 4d; Supplementary Movie 1). This behaviour indicates that AMBRA1 is a diffusible nuclear protein rather than stably anchored to chromatin.

### AMBRA1 alters transcriptional responses to light and darkness

To investigate the transcriptional changes associated with loss of AMBRA1, we performed RNA sequencing of WT and *ambra1a/b* plants grown under long-day conditions or in continuous darkness (Figs. 2a and 3a). Principal component analysis separated samples primarily according to light regime along PC1, which explained 39.4% of the variance, and according to genotype along PC2, which explained 14.3% of the variance (Supplementary Fig. 11). We identified 1,023 differentially expressed genes in *ambra1a/b* relative to WT under long-day conditions, of which 654 (64%) were upregulated, and 1,165 in continuous darkness, of which 706 (60.6%) were downregulated (Supplementary Fig. 11 and Tables S4 and S5).

Gene Ontology and KEGG enrichment analyses identified distinct functional signatures under the two growth conditions (Fig. 5a and 5b and Table S6). Under long-day conditions, genes associated with amino acid and carbon catabolism were enriched among the upregulated genes, whereas genes related to photosynthesis (Fig. 5a, Supplementary Fig. 12, Supplementary Table 6) were enriched among the downregulated genes (Fig. 5a and Supplementary Table 6). In darkness, photosynthesis-associated genes were enriched among the upregulated genes, while genes involved in carbon and nitrogen assimilation were predominantly downregulated (Fig. 5b, Supplementary Fig. 12, Supplementary Table 6). The accumulation of photosynthesis-associated transcripts in dark-grown *ambra1a/b* plants is consistent with their increased chlorophyll level and impaired skotonemal development, supporting a requirement for AMBRA1 to repress photomorphogenesis in darkness. We next examined the 317 genes differentially expressed in both conditions (Fig. 5c and Supplementary Table 7). Of these, 283 showed the same direction of change under long-day conditions and in darkness, whereas 34 displayed opposite responses (Fig. 5d and Supplementary Table 8). Downregulated across both regimes were key cell wall biosynthesis and extension networks, including cellulose synthase complexes, xyloglucan endotransglycosylases, pectin lyases, and expansins (Supplementary Fig. 12). This suppression was coordinated with the downregulation of essential cell cycle and mitotic machinery components, including A- and B-type cyclin-dependent kinases, cyclin B, cyclin D, phragmoplast-associated kinesins, and core histones. These expression changes are consistent with the reduced cell dimensions and impaired colony growth of the mutant. Loss of AMBRA1 was also accompanied by changes in stress- and hormone-responsive gene expression. Under both conditions, *ambra1a/b* plants showed increased abundance of transcripts encoding components of abscisic acid signalling and several late embryogenesis abundant (LEA) proteins. Transcripts associated the 12-oxo-phytodienoic acid (OPDA) biosynthetic pathway (e.g. *LOX*, *AOS*, *AOC*) and OPDA-responsive signalling were also more abundant (Supplementary Fig. 12, Supplementary Table 7).

**Figure 5.**
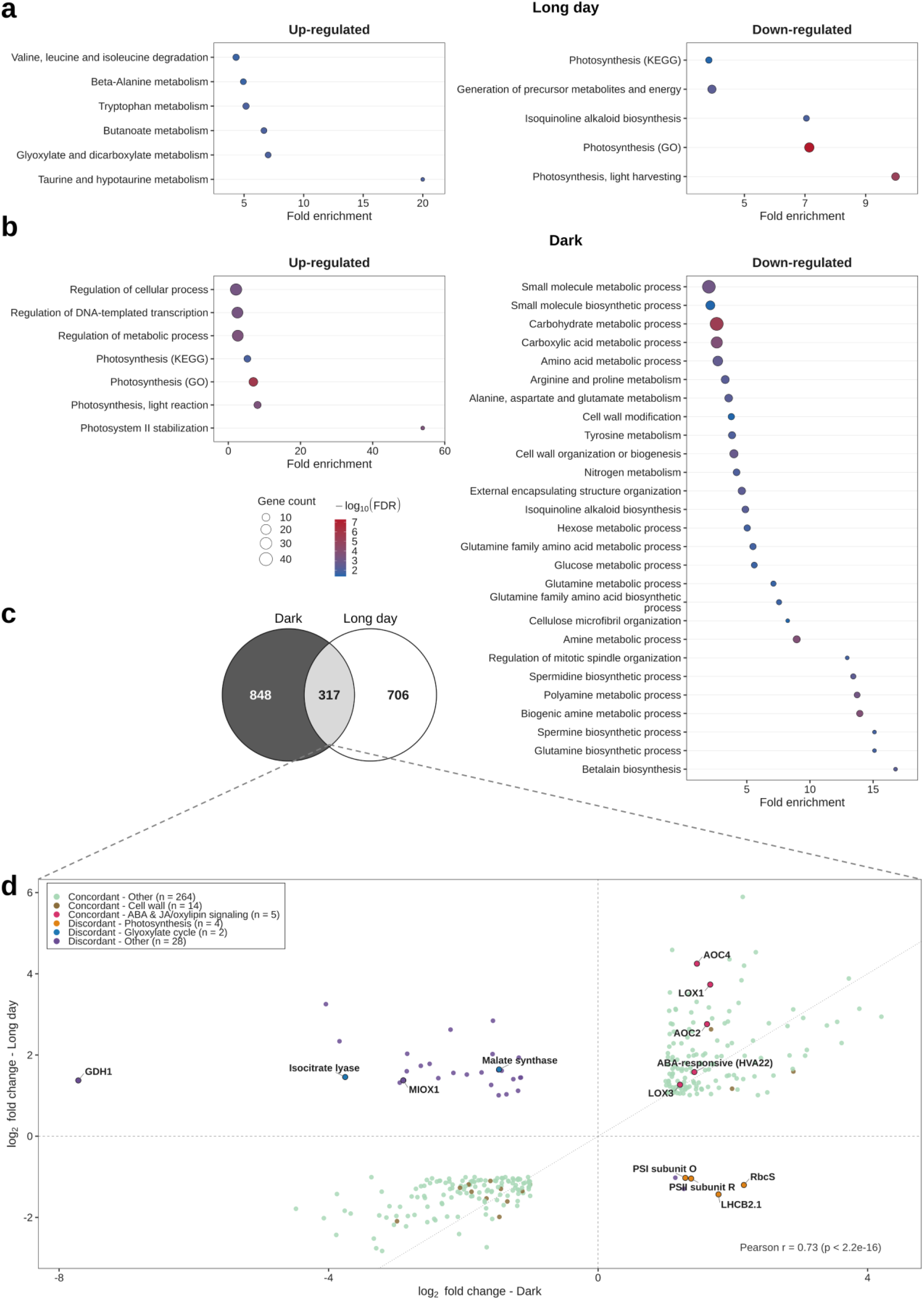
Transcriptome profile of WT and *ambra1a/b* plants grown under a long-day photoperiod or in continuous darkness. a-b) Functional enrichment analysis (GO terms and KEGG pathways, FDR < 0.05) of genes upregulated and downregulated in *ambra1a/b* vs WT under long-day conditions (a), and upregulated and downregulated under dark conditions (b). For all dot plots, the x-axis represents Fold Enrichment, dot size represents the number of genes (Gene Count) enriched in each term, and the color gradient reflects statistical significance (−log10 FDR), from blue (lower significance) to red (higher significance). c) Venn diagram showing the overlap of DEGs between long-day and dark conditions. d) Scatter plot comparing the log2 fold change of the 317 DEGs common to both conditions (log2FC Dark on the x-axis, log2FC long day on the y-axis); concordantly (green, n = 283, 89%) and discordantly (purple, n = 34, 11%) regulated genes are indicated, genes belonging to specific functional categories (cell wall, ABA & JA/oxylipin signaling, photosynthesis, glyoxylate cycle) are colored and counted as indicated in the legend. Pearson r = 0.73 (p < 2.2 × 10⁻¹⁶). Specific genes are labeled by name as in the legend.

Among the genes showing contrasting expression patterns in light and darkness were several associated with primary metabolism, including those encoding isocitrate lyase, malate synthase, GDH1 and MIOX1. Notably, this group also included several photosynthesis-associated genes. For instance, transcripts encoding the small subunit of ribulose-1,5-bisphosphate carboxylase/oxygenase and photosystem subunits were reduced in *ambra1a/b* under long-day conditions but increased in darkness (Fig. 5d and Supplementary Table 8).

### AMBRA1 and COP1 functions partially converge in darkness

COP1 is a key repressor of photomorphogenesis in plants, and a mutant deficient in all nine COP1 homologs in *P. patens* (*cop1_9x* KO) also shows partial derepression of photomorphogenesis in the dark. Given the strong photomorphogenic phenotype of *ambra1a/b* plants in the dark, we compared their growth with the *cop1_9x* KO mutant^23^ (Fig. 6).

**Figure 6.**
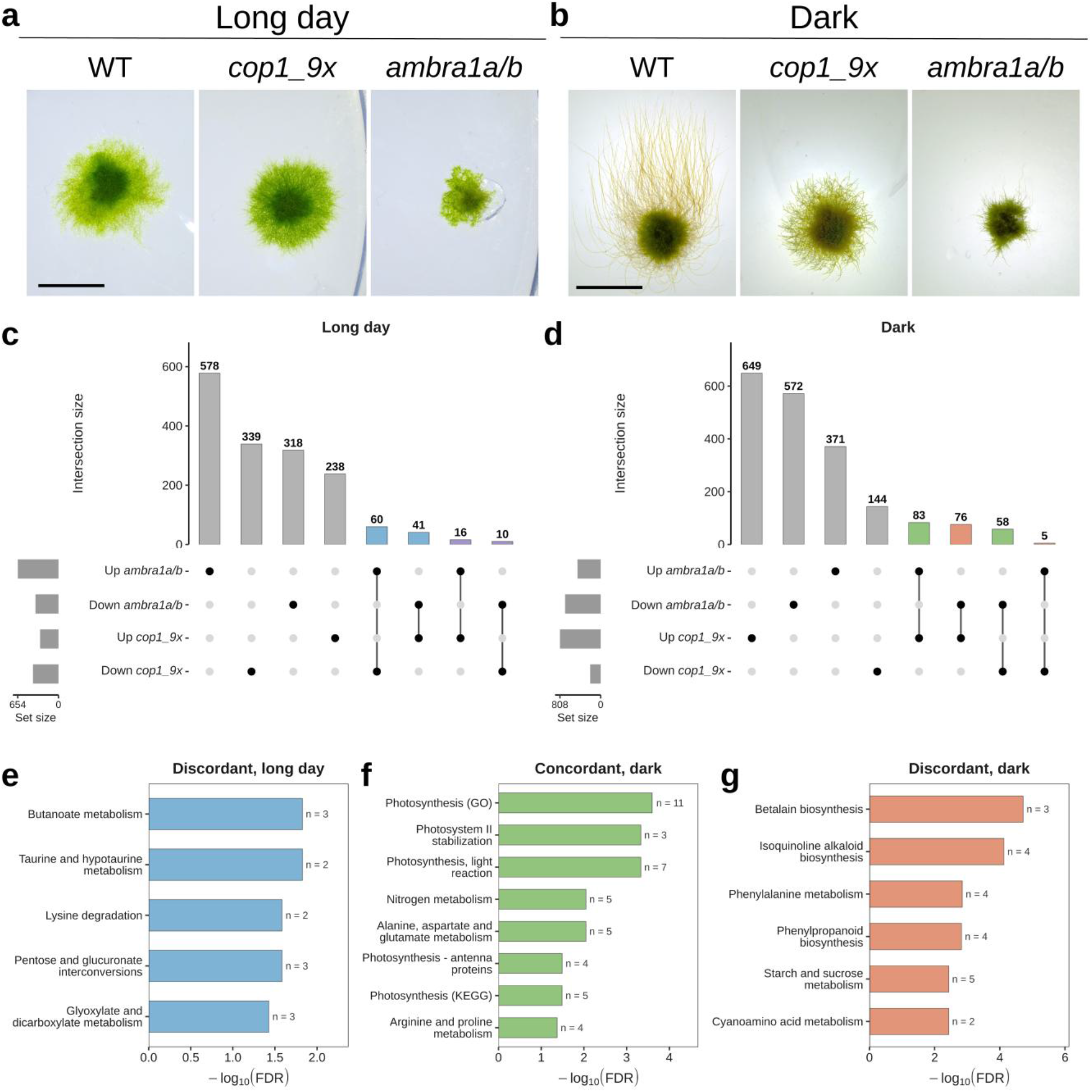
Comparative phenotypic and transcriptomic analysis of *ambra1a/b* and *cop1_9x* mutants under long-day and dark conditions. a-b) Representative images of WT, *cop1_9x*, and *ambra1a/b* colonies grown under long-day (a) and dark (b) conditions. Scale bar = 5 mm. c-d) Upset Plot showing the overlap of differentially expressed genes (DEGs) between *ambra1a/b* vs WT and *cop1_9x* vs WT under long-day (c) and dark (d) conditions. e-g) Gene Ontology (biological process) enrichment and KEGG analysis for the discordant DEGs shared between *ambra1a/b* and *cop1_9x* under long day (e), concordant (f) and discordant DEGs shared between *ambra1a/b* and *cop1_9x* in the dark; the x-axis shows −log10(FDR) and gene counts for each term are indicated next to the bars (FDR < 0.05).

Under long day conditions, both WT and *cop1_9x* KO lines displayed characteristic radial filamentous growth, but the *ambra1a/b* mutant exhibited reduced growth (Fig. 6a). In continuous darkness, WT successfully initiated the classic skotomorphogenic program by extending etiolated caulonemal filaments. The *cop1_9x* mutant continued to grow but failed to form gravitropically responsive filaments, consistent with constitutive photomorphogenesis. The *ambra1a/b* mutant produced few filaments, which failed to elongate, and remained substantially smaller than the other genotypes (Fig. 6b). These phenotypes indicate that AMBRA1 and COP1 both contribute to the repression of photomorphogenesis in darkness, although their loss has distinct effects on protonema growth and development. We therefore asked whether this partial phenotypic convergence was also reflected at the transcriptional level, and compared our *ambra1a/b* RNA-sequencing dataset with previously published transcriptomic data from the *cop1_9x* mutant^23^ (Fig. 6c-d). Most genes differentially expressed in *ambra1a/b* were not differentially expressed in *cop1_9x*. Under light conditions, the two datasets shared 127 differentially expressed genes, corresponding to 12.4% of the genes differentially expressed in *ambra1a/b* (Fig. 6c and Supplementary Table 9). In continuous darkness, 222 genes were shared, corresponding to 19.1% of the *ambra1a/b* differentially expressed genes (Fig. 6d and Supplementary Table 10).

Under light conditions, 80% of the shared genes changed in opposite directions in the two mutants. Functional enrichment analysis associated these genes primarily with carbohydrate and amino acid metabolism (Fig. 6e and Supplementary Table 11). In continuous darkness, by contrast, 64% of the shared genes changed in the same direction. Gene Ontology and KEGG enrichment analyses showed that these concordantly altered genes were enriched in photosynthesis-associated processes and pathways related to precursor and amino acid metabolism (Fig. 6f and Supplementary Table 11). This shared set included genes encoding photosystem I and II components and the small subunit of ribulose-1,5-bisphosphate carboxylase/oxygenase, whose transcripts were more abundant in both mutants than in WT. Genes changing in opposite directions in darkness were instead enriched in phenylpropanoid, carbohydrate and amino acid metabolism (Fig. 6g and Supplementary Table 11).

Together, the distinct developmental phenotypes and limited transcriptional overlap indicate that AMBRA1 and COP1 have largely distinct functions. Nevertheless, the concordant accumulation of photosynthesis-associated transcripts in darkness suggests that both proteins contribute to the repression of photosynthetic development in the absence of light.

## Discussion

Light signalling represents a master regulatory system that translates environmental cues into developmental responses to optimize plant growth. Extensive research has identified E3 ubiquitin ligase complexes, with the COP1 E3 ubiquitin ligase playing a central role in repressing photomorphogenesis in the dark by targeting positive regulators of light signalling for degradation^5,24^. Although this pathway has been extensively characterized, the regulatory network coordinating developmental responses to changing light conditions remains incomplete. Here, we identify AMBRA1 as a previously unrecognized nuclear WD40 protein that integrates light availability with developmental programs. While mammalian AMBRA1 has been studied as a regulator of autophagy, cell proliferation and ubiquitin-mediated proteolysis^17,19^, its biological function in plants has remained entirely unknown.

Our bioinformatic and genomic analyses reveal that AMBRA1 is deeply conserved across the photosynthetic eukaryotes and traces back to a single origin prior to land plant evolution. AMBRA1 copy number is tightly regulated across kingdoms. While lineage-specific whole-genome duplications (WGDs) led to the retention of multiple *AMBRA1* paralogs in several plant taxa, other lineages have a single copy. Similarly, most vertebrate lineages maintain a single *AMBRA1* gene copy despite early rounds of WGDs. The presence of two copies in bony fishes is an exception attributed to their lineage-specific third round of WGDs^12,13^. This cross-kingdom tendency to constrain *AMBRA1* duplication suggests strong evolutionary pressure to maintain precise gene dosage.

Structural modelling supports the evolutionary conservation of AMBRA1. The predicted *P. patens* protein retains the characteristic seven-bladed WD40 β-propeller that forms the structural core of mammalian AMBRA1. In both kingdoms, rather than being joined by a standard three-residue loop, the WD40-N and WD40-C subdomains are separated by extensive intrinsically disordered region (407 and 565 residues in *P. patens* and human AMBRA1, respectively). In *A. thaliana*, AtAMBRA1a (At1g04140) is among the WD40 proteins bearing WDxR motifs catalogued as candidate DDB1-binding partners, yet no interaction was detected in yeast two-hybrid assays^8^. AMBRA1 WD40 domain is predicted to fold only upon binding DDB1^16^; however this binding-induced folding may not be efficiently coordinated by the yeast two-hybrid system^25^. In tomato instead, a predicted protein-protein interaction network integrating orthology, domain architecture, coexpression, and information transferred from model organisms, places the AMBRA1a ortholog (Solyc05g053130.2) among the predicted partners of DDB1^26^.

The developmental phenotypes of the *P. patens ambra1a/b* mutant establish AMBRA1 as a central regulator of plant growth rather than a factor acting on a single developmental process. Under illumination, AMBRA1 is determinant for i) cell fate, regulating the normal transition between the two filament types that constitute the protonema, chloronema and caulonema, ii) cell size, iii) plant growth, iv) gravitropic response.

In mosses, protonemal development integrates hormonal, metabolic and environmental cues, with ABA acting as a negative regulator of chloronema-to-caulonema progression by antagonizing auxin signalling and limiting cyclin expression and cellular proliferation^27–30^. The *ambra1a/b* transcriptome consistently reveals an activation of ABA- and OPDA-responsive signatures, particularly under light conditions. Since ABA biosynthetic genes are not affected, ABA-responsive signatures may reflect hormone perception or downstream signalling rather than hormone production. Similarly, OPDA, which in *P. patens* functions as a bioactive signalling molecule in the absence of canonical downstream jasmonic acid, has been shown to inhibit growth, consistent with the reduced growth of *ambra1a/b* mutants^31^. Consistent with this hormonal reprogramming, *ambra1a/b* mutants show reduced cell-cycle activity. This suggests a conserved role of AMBRA1 in cell-cycle regulation across animals and plants^15^, though with opposite directional effects: while mammalian AMBRA1 deficiency causes uncontrolled proliferation, plant mutants display reduced cell-cycle activity. The identification of *Arabidopsis* AMBRA1a/b as putative E2F targets suggests that also the association between AMBRA1 and proliferation could be evolutionarily conserved^32^.

Under prolonged darkness, AMBRA1 is required to establish the dark developmental state, including skotonemata formation and gravitropic filament elongation, and to repress photosynthesis-associated genes at both the transcript and protein levels, as well as glyoxylate cycle genes. These findings indicate a role for AMBRA1 in repressing photomorphogenesis. This role is reminiscent of COP1, a well characterised repressor of photomorphogenesis, which, like AMBRA1, is conserved across kingdoms and can form E3 ligase complexes through its WD40 domain^5,7^. Comparative physiological and transcriptomic analyses indicate that AMBRA1 and COP1 converge on the repression of photomorphogenesis but regulate distinct pathways^23^. Under light conditions, *ambra1* mutants form smaller colonies than *cop1_9x* mutant. In the dark, both mutants fail to develop skotonemata, and *ambra1* colonies are even smaller. This partial phenotypic overlap is consistent with the transcriptomic data: neither mutant represses photosynthesis-associated genes in the dark, whereas loss of AMBRA1 uniquely affects genes associated with cell cycle and cell wall biogenesis in darkness, and with photosynthesis and carbohydrate metabolism under long-day conditions. Although mammalian AMBRA1 is a core regulator of autophagy initiation via Beclin1-PI3KC3, we found no evidence of altered expression of core ATG genes in *ambra1a/b* mutants, suggesting that this function may not be conserved in plants, or that it operates independently of transcriptional regulation.

AMBRA1 was observed in the nucleus in plants maintained under long-day conditions or prolonged darkness. Within the nucleus, AMBRA1 formed punctate speckles, a pattern observed in some proteins with intrinsically disordered regions and suggestive of an additional level of spatial organization^33^. AMBRA1 speckles resemble photo-bodies, speckles formed by phytochromes and many other light signalling components^34^. Whether AMBRA1 speckles have a similar role remains to be determined.

Overall, our work identifies AMBRA1 as new integrator of light availability and plant development, highlighting an essential role for repression of photomorphogenic traits in darkness. Defining its direct molecular partners across these pathways will be essential to understand how a single evolutionarily conserved scaffold orchestrates such diverse developmental outputs in plants.

## Methods

### Phylogenetic analyses

To reconstruct the phylogeny of AMBRA1, homologous protein sequences to *Homo sapiens* HsAMBRA1 (NP_001373940.1) were retrieved from the NCBI database and the Joint Genome Institute (JGI) Plant Comparative Genomics portal Phytozome. In total, 92 AMBRA1 sequences from diverse animals, plants, and algal species were collected (Supplementary Table 1), together with 14 KATNB1 and 9 WDR5 sequences as outgroups (n = 23), yielding 115 sequences for alignment. When a species contained two or more AMBRA1 genes, the paralogs were designated a, b (and c) according to their position in the genome (in order of chromosome or scaffold and coordinate), because the phylogeny did not show a clear separation of the paralogs into distinct subfamilies. Protein sequences were aligned using MAFFT v7.505 with the E-INS-i strategy (--genafpair --maxiterate 1000). The resulting alignment comprised 115 sequences and 4,581 amino-acid positions. Conserved regions suitable for phylogenetic analysis were selected with the Gblocks algorithm using relaxed parameters as in^12^. Maximum-likelihood phylogenetic inference was performed with IQ-TREE v2.0.7; the best-fit substitution model (LG+R5) was selected by ModelFinder according to the Bayesian information criterion. Branch support was assessed with 1,000 ultrafast bootstrap replicates, with nearest-neighbour-interchange optimization of bootstrap trees (-bnni), and with 1,000 SH-like approximate likelihood ratio test replicates. Phylogenetic inference was conducted on an unrooted tree; for visualization, the tree was rooted on the monophyletic KATNB1 clade using the root function of the R package ape, which did not alter the inferred topology. The tree was drawn as a cladogram (branch lengths not to scale) in R with ape and ggplot2.

### AMBRA1 WD40 alignment

To define homologous WD40 regions consistently across the dataset, PpAMBRA1a was used as the reference sequence for both the N- and C-terminal analyses. InterProScan identified an N-terminal WD40/YVTN repeat-like domain in PpAMBRA1a spanning residues 113-343. For the N-terminal analysis, residues 103-353 were selected to include 10 flanking amino acids on each side of the predicted domain. The region selected for the C-terminal analysis comprised residues 711-917 of PpAMBRA1a and included the C-terminal WD40 repeat identified by SMART at residues 830-870. The columns corresponding to the two PpAMBRA1a reference intervals were located in the full-length MAFFT alignment and projected onto each of the 92 AMBRA1 proteins. The projected N- and C-terminal regions were then realigned independently using MAFFT v7.505 with the E-INS-i strategy (--genafpair --maxiterate 1000). For visualization, only alignment columns containing a residue (i.e. not a gap) in at least 50% of the sequences were retained. After filtering, the N-terminal alignment comprised 247 positions and the C-terminal alignment comprised 186 positions. For each retained alignment position, the consensus amino acid was defined as the most frequent residue after excluding gaps. Conservation was calculated as the proportion of non-gap, matching the consensus amino acid at that position. Residues matching the consensus were shaded dark grey at positions with at least 70% conservation and light grey at positions with 50-69% conservation. Residues that did not match the consensus and positions with less than 50% conservation were left unshaded. The N- and C-terminal regions were displayed separately as shaded conservation maps, with sequences in the same order as in the phylogenetic tree (Supplementary Fig. 1).

### AMBRA1 structural predictions

Structural models of PpAMBRA1a (AlphaFold ID: A0A2K1JYD0-F1), PpAMBRA1b (AlphaFold ID: A0A2K1JGJ8-F1), and HsAMBRA1 (AlphaFold ID: Q9C0C7-4-F1) were retrieved from the AlphaFold Protein Structure Database^35^ and visualized using PyMOL. The WD40 core domains were manually annotated according to previous work on HsAMBRA1 ^16^.

### Expression profile of P. patens genes during dark-to-light transition

To retrieve and analyze *PpAMBRA1a* (Pp3c10_9370) and *PpAMBRA1b* (Pp3c14_5930) and light signaling^9^ gene expression, we used a previously collected dataset from our laboratory^21^. Raw transcript expression values (estimated counts from transcript-level quantification) for previously published WT *P. patens* samples across three time points in 12h light/ 12h dark photoperiod (0, 2, and 6 hours of light exposure, time 0 corresponded to the end of the night) were rounded to the nearest integer to satisfy the integer-count requirement of downstream statistical analysis. Data handling, preprocessing, and normalization were conducted within the R statistical environment using the DESeq2 framework. Low-abundance transcripts were filtered out by retaining only genes with non-zero counts in at least four samples. Finally, count data were subjected to a variance-stabilizing transformation (VST) to produce normalized expression values for downstream analyses. Diurnal expression profiles of *AtAMBRA1a* (At1g04140) and *AtAMBRA1b* (At5g43930) (Supplementary Fig. 4) were retrieved from the DIURNAL database (http://diurnal.mocklerlab.org/) using the COL_LDHH dataset^36^.

### Correlation matrix

Pairwise gene-gene expression correlations were computed using Pearson correlation coefficients based on normalized expression values. A curated set of photomorphogenesis-related genes^9^ was used for the analysis. Correlation matrices were generated using pairwise complete observations. Hierarchical clustering of genes was performed using Euclidean distance and complete linkage based on correlation profiles. The resulting correlation matrix was visualized as a heatmap using the pheatmap package in R. Branches were manually rotated to visualize PpAMBRA1a and PpAMBRA1b on top, without altering any cluster assignment.

### Plasmid constructs used to generate ambra1 mutant lines

Regions from the *AMBRA1a* and *AMBRA1b* genes were amplified by PCR from WT genomic DNA (gDNA) and cloned up- and down-stream of the coding sequence of the hygromycin-B and G418 resistance cassettes (see Supplementary Fig. 6). The left homologous arm of *AMBRA1a* was amplified via PCR using the primers P1_AMBRA1a and P2_AMBRA1a and was then digested and inserted into the AvrII and XhoI sites of the BHRf plasmid ^37^. The right homologous arm was amplified using the primers P3_AMBRA1a and P4_AMBRA1a and was then digested and inserted into the SpeI and AscI sites of the former plasmid to make the AMBRA1a-BHRF plasmid.

The left homologous arm of AMBRA1b was amplified via PCR using the primers P1-Am1b_up_for and P2-Am1b_up_rev, and was then digested and inserted into the AvrII and XhoI sites of the BNRf plasmid ^37^. The right homologous arm was amplified using primers P3-Am1b_up_for and P4-Am1b_up_rev and was then digested and inserted into the SpeI and AscI sites of the former plasmid to make the AMBRA1b-BNRF plasmid. Primers list is in Supplementary Table 12.

For constitutive expression of the full-length coding sequence, *AMBRA1a* (Pp3c10_9370V3.3) and *AMBRA1b* (Pp3c14_5930V3.1) were PCR amplified from cDNA with the attB-AMBRA1aF / attB-AMBRA1aR primers and attB-AMBRA1F / attB-AMBRA1R primers (Primers list is in Supplementary Table 12), subcloned in pDNR201 via Gateway BP reaction and then transferred via a Gateway LR reaction into the pT1OGY vector under control of PpEF1 promoter and in frame with the YFP sequence at 3’ end, before the stop codon (https://www.nibb.ac.jp/evodevo/PHYSCOmanual/11.6_revised_170727.htm).

### Transformation of P. patens

*P. patens* transformation was performed as described in ^37^ with minor modifications. Multiple independent transformations were conducted to generate the *ambra1a*, *ambra1b*, *ambra1a/b*, and *ambra1a/b* (Cre) mutant lines, as well as the reporter lines expressing AMBRA1A-YFP or AMBRA1B-YFP.

Protoplasts were isolated from 6-day old protonema tissue cultured on PPNH_4_ medium with 0.72 % (w/v) agar. Protoplasts were transformed by polyethylene glycol (PEG)-mediated heat shock with 20 μg of the plasmid DNA linearized with restriction enzymes (AvrII and AscI for both *ambra1a* and *ambra1b* KO; PmeI for AMBRA1a and AMBRA1b overexpressors). After transformation, protoplasts were incubated in liquid PpNH_4_, 6.6 % (w/v) D-mannitol for 24 h in the dark. An equal volume of 8.5% (w/v) D-mannitol and 14% agar was added before plating protoplasts on PpNH4, 6.6% (w/v) D - mannitol overlaid with cellophane. Plates were incubated in standard growth conditions for 5 days. Regenerated protoplasts were first transferred to PpNH4 medium with 1% (w/v) agar and the appropriate antibiotic (25 ng/µl hygromycin, 50 ng/µl G418, or 50 ng/µl zeocin, depending on the transformation) for another 10-14 days and then to normal PpNH_4_ medium with 0.72 % (w/v) agar for 2 weeks. After another round of selection on antibiotics, stable resistant lines were grown for 2 weeks in standard growth conditions.

Genomic DNA was then extracted from each clone and mutation of the target genes tested by PCR and sequencing (Supplementary Fig. 6). We kept at least two independent lines for each target gene to address the *in planta* function of *PpAMBRA1*. The four single-KO lines (two independent *ambra1a* and two independent *ambra1b* lines) lacked *PpAMBRA1a* and *PpAMBRA1b* transcripts respectively (Supplementary Fig. 6). By the same strategy, we isolated three independent double mutants *ambra1a/b* KO by knocking out *PpAMBRA1b* in an *ambra1a* #1 KO moss (Supplementary Fig. 6). The three *ambra1a/b* KO lines lacked both *PpAMBRA1a* and *PpAMBRA1b* transcripts (Supplementary Fig. 6). This was confirmed using two sets of primers specific for the two genes but also by using primers that amplify simultaneously *PpAMBRA1a* and *PpAMBRA1b* (Supplementary Fig. 6d). The presence of flanking loxP sites (Supplementary Fig. 8) was utilized to excise the resistance cassette via transient expression of Cre recombinase in the *ambra1a/b* double KO background, resulting in a marker-free deletion of the target genes ^38^. The resulting mutant lines were verified by PCR amplification (Supplementary Fig. 8) and the genomic architecture was further confirmed by Sanger sequencing of the entire recombinant loci.

For constitutive overexpression of YFP-containing AMBRA1 coding sequences isolated clones were genotyped with primers targeting the insertion site and screened for YFP expression using inverted fluorescent microscope (Leica DMI4000B) with standard GFP filter set. Confirmed clones were grown vertically in the dark on PpNH₄ medium for two weeks to assess functional complementation of mutant phenotype by the introduced constructs.

### Plant material, growth conditions and phenotypic analysis

The Gransden (Gd) WT, *ambra1a* KO, *ambra1b* KO, *ambra1a/b* KO, *AMBRA1a-YFP*, *AMBRA1b-YFP* and *cop1_9x* KO ^23^ mutant strains of *P. patens* were either cultivated on PPNO_3_ or PPNH_4_ medium with 0.72 % (w/v) agar ^37,39^ in a growth chamber at 24°C under a 16 h : 8 h, light : dark long day photoperiod. For light growth tests, protonemata were pre-cultivated for 9 days on PpNO_3_ medium, and 2-mm spots were transferred to fresh PpNO_3_ plates and grown for 28 days under standard conditions. For dark growth tests, protonemata pre-cultivated for 7 days on PpNH_4_ were transferred as 2-mm spots onto PPNH_4_ plates and grown horizontally for 7 days under standard conditions, after which plates were wrapped in aluminium foil and incubated vertically in the dark for 14 days at 22°C. Colonies were imaged with a Leica Ivesta 3 stereomicroscope. Colony area (light-grown colonies) was quantified in Fiji. For dark-grown colonies, skotonemata density was quantified as the integrated optical density of a 1-mm-high strip just above the colony margin (image thresholded to 8-bit with manual background subtraction), and skotonema length as the mean length of the six longest skotonemal filaments per colony, both in Fiji.

### Calcofluor staining and cell morphometrics

The chloronema-to-caulonema developmental transition was quantified on *P. patens* protonemal tissue cultivated on PPNO_3_ medium for 7 days under standard light conditions (16-h light/8-h dark). To grow the colonies at low density, one-week-old protonemata were ground for 15 sec., sieved in a 100 µm mesh, and grown for a week. Prior to imaging, the tissue was stained with Calcofluor White (10 µg/ml dye solution) to delineate cell boundaries and imaged with a Leica DMI6000 B inverted fluorescence microscope. To avoid mis-classification of still-dividing cells, measurements were restricted to subapical cells. Cell length (longitudinal axis, from the centre of the proximal to the centre of the distal cross wall) and width (diameter at the cell midpoint) were measured in Fiji (ImageJ); at least 300 subapical cells per genotype were measured. Raw measurements were imported into R (v4.5.2), reshaped to long format and annotated by filament position (first/second/third subapical or branch cell) and genotype; branch cells and unidentified measurements were excluded (dplyr). For each cell type (chloronema/caulonema), genotype and filament position, mean length and width, SEM and 95% confidence intervals were calculated, and data were visualized in ggplot2 as jittered points with mean crossbars and SEM/95% CI error bars.

### Confocal microscopy

Stable lines expressing AMBRA1a-YFP or AMBRA1b-YFP were grown on PPNH_4_ or PpNO_3_ medium for 7 days under continuous light at 50 µmol photons m⁻² s⁻¹ (light-grown samples) or for 7 days in light followed by 14 days in darkness, as described for the growth test (dark-grown samples). For detailed analysis, a 63× oil-immersion objective HC PL APO CS2 63×/1.40 OIL) was used. Images were acquired with two channels: YFP (excitation 514 nm, emission 520-575 nm) and chlorophyll *a* (excitation 478 nm, emission 650-750 nm). Z-stacks (1-µm step, between 7 and 13 planes) were acquired. Further processing in Fiji included deconvolution (Iterative Deconvolve 3D), chloroplast bleed-through subtraction, maximum-intensity projections, and brightness and contrast adjustments. For cell division analysis, the protonemata were cultured in a microfluidic device (PDMS) for 24 hr before imaging.

### Transcriptome analysis

Total RNA was extracted from protonemata (light- and dark-grown colonies) using the FastPure Universal Plant Total RNA Isolation Kit (Vazyme). 3′ mRNA-seq libraries (QuantSeq 3′ mRNA-Seq V2 FWD, Lexogen) were sequenced on an Illumina NextSeq 500 (single-end, 150 nt). For each condition, two independent *ambra1a/b* KO clones were used for the analyses: one retaining the resistance cassettes used for gene targeting, and one in which these cassettes were removed via Cre-lox-mediated excision, in order to distinguish transcriptional changes attributable to the ambra1a/b mutation from cassette-specific effects. Illumina adaptors and poly(A) tails were trimmed with cutadapt v4.4, and reads were aligned to the *P. patens* v3.3 transcriptome (Phytozome) with STAR v2.7.11b and quantified with Salmon v1.10.2. Counts per million (CPM) were derived from these quantifications, and, after removing genes with CPM ≤ 1 in fewer than 3 samples, log2-transformed CPM values were used for principal component analysis (PCA; prcomp, unit-variance scaling) to assess sample clustering by genotype. Three-to-four biological replicates per genotype were initially generated; following PCA-based outlier exclusion, three-to-four biological replicates were retained for WT in each condition, while combining both *ambra1a/b* clones (cassette-retaining and cassette-excised) yielded five-to-six replicates per condition for the KO group. As the two independent *ambra1a/b* clones showed comparable transcriptional profiles, they were grouped together for downstream comparisons against WT. For differential expression, counts were normalized with the trimmed mean of M-values (TMM) method and compared between WT and *ambra1a/b* with edgeR (v4.6.2, negative binomial model); genes with FDR < 0.05 and |log₂FC| > 1 were called differentially expressed (DEGs).

Gene Ontology (GO) annotations for *P. patens* were retrieved from the Phytozome v3.3 annotation file (locusName-GO mapping) and restricted to the Biological Process (BP) domain. To account for the hierarchical structure of the GO, each gene’s direct BP annotations were propagated to all ancestor terms up to the root, using the GOBPANCESTOR mapping from the Bioconductor package GO.db. For each list of differentially expressed genes (DEGs; FDR < 0.05, |log2FC| > 1), enrichment of each GO BP term was tested using a one-sided hypergeometric test against a background set consisting of all genes detected in the corresponding expression dataset (CPM table) that carried at least one BP annotation. Fold enrichment was calculated as the ratio between the proportion of DEGs annotated with a given term and the proportion of background genes annotated with the same term. Resulting p-values were corrected for multiple testing using the Benjamini-Hochberg (BH) procedure, and terms with FDR < 0.05 were considered significantly enriched.

For KEGG pathway enrichment analysis, Genes were first linked to KEGG Orthology (KO) identifiers using the same Phytozome v3.3 annotation file, and KOs were mapped to KEGG pathways using a static KO-pathway reference table derived from PICRUSt2, from which human-disease-specific pathways were excluded as not relevant to a moss species. Enrichment testing followed the same approach used for GO terms: a one-sided hypergeometric test against a background of KO-annotated genes within the same background set used for the corresponding GO analysis, with Benjamini-Hochberg correction and a significance threshold of FDR < 0.05. Significant GO BP and KEGG pathway terms (FDR < 0.05) were combined into single dot plots (fold enrichment on the x-axis, dot size proportional to the number of genes, color scaled to -log10(FDR)). Because GO and KEGG occasionally described the same underlying biological process under different names (e.g., “photosynthesis, light harvesting in photosystem I” vs. “Photosynthesis - antenna proteins”), pairs of significant terms sharing ≥90% of their annotated genes (Jaccard index) were merged into a single point, retaining the gene union and the fold enrichment/FDR of the more significant of the two.

### Comparative transcriptome with cop1_9x KO

AMBRA1 DEG lists (dark and light, *ambra1a/b* vs WT) were compared with the published *cop1_9x* RNA-seq dataset of Kreiss et al. ^23^, restricting the comparison to mutant-vs-WT within the same light condition (dark with dark, light with long day). For genes differentially expressed in both datasets, concordance of regulation direction was evaluated by Pearson correlation between the AMBRA1 log_2_ FC and the Kreiss M-value. GO enrichment and KEGG analysis was performed as above.

### Statistical analysis

All quantitative data were visualized using the ggplot2 package in R. For comparisons between two groups (colony area at each timepoint; pigment content metrics, chlorophyll/fresh weight, chlorophyll a/b ratio, chlorophyll/carotenoid ratio, in WT vs *ambra1a/b*), normality was first assessed for each group with the Shapiro-Wilk test: if both groups were normally distributed, homogeneity of variance was tested with an F-test and a Student’s or Welch’s t-test was applied accordingly; otherwise, a Wilcoxon rank-sum test was used. For comparisons among the four genotypes (WT, *ambra1a*, *ambra1b*, *ambra1a/b*; fresh weight, skotonemata density, skotonemata length), an omnibus Kruskal-Wallis test was followed by pairwise Dunn’s post-hoc tests with Benjamini-Hochberg correction across all genotype pairs, and results are summarized as a compact letter display (groups sharing a letter are not significantly different at p < 0.05). Significance in two-group comparisons is indicated with asterisks (*p < 0.05, **p < 0.01, ***p < 0.001, ****p < 0.0001).

## Author contributions

CB, VA and LV contributed to the investigation of the study; CB and AA contributed to the writing - original draft of the study; CB, LDV, AH, TM and AA contributed to the writing - review & editing of the study. AA and LDV contributed to the supervision of the study. LDV, TM and AA contributed to the conceptualization of the study.

## Acknowledgment

The authors thank Prof. Ute Hoecker for *cop1* line, Anna Segalla, Alessandra Bellan, Jessica Ceccato and Ilena Zuin for mutant isolation and preliminary characterization of null mutant lines. AH was supported by the German Research Foundation (DFG) under Germany’s Excellence Strategy (EXC-2189 - Project ID 390939984) and in the framework of MAdLand (https://madland.science, DFG priority programme 2237; HI 1369/8-2, Project ID 440515704). LV was partially supported by US-NSF (MCB 2154573 and 2439582). AI was used to assist with figure assembly without altering the image data, and to revise the English language of the manuscript without changing its scientific content.

## Data Availability

The RNA-seq data generated in this study have been deposited in the NCBI Gene Expression Omnibus (GEO) database under accession code GSE350707. The raw data used in this study have been deposited in Research Data Unipd and are available at https://researchdata.cab.unipd.it/1937/.

