## Supplemental Figures for "Evolutionary co-option of the ancient protein AMBRA1 in plant photomorphogenesis"

### Supplementary Figures 1-12

#### List of Supplementary Tables and Movie

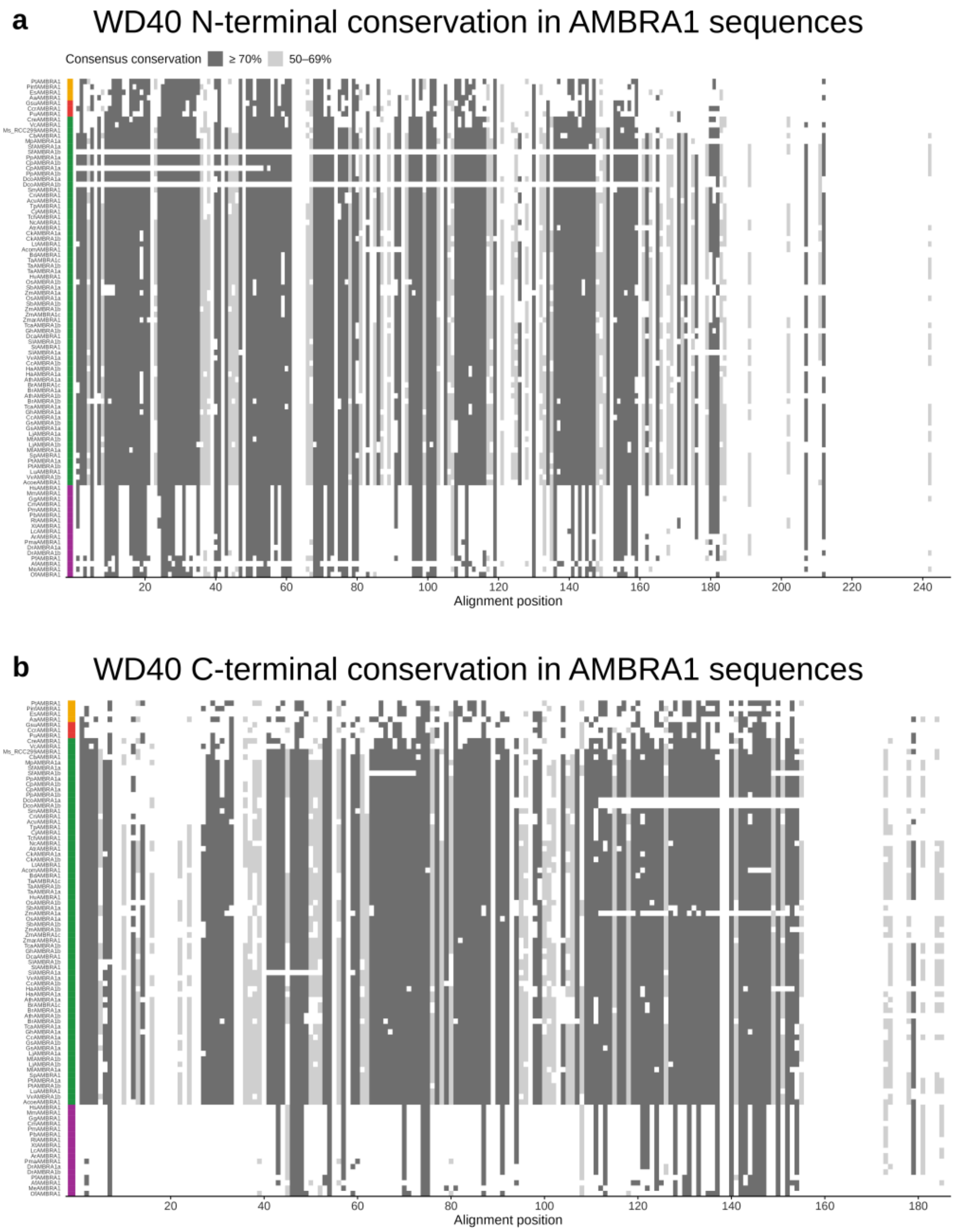


**Supplementary Fig. 1. Sequence conservation of AMBRA1 N-terminal (a) and C-terminal (b) WD40 domains across 92 AMBRA1 homologues from Metazoa, Stramenopiles, Rhodophyta and Viridiplantae**. a) Multiple-sequence alignment of regions homologous to the PpAMBRA1a N-terminal WD40 domain (reference residues 103-353) in 92 AMBRA1 proteins. b) Multiple-sequence alignment of regions homologous to the PpAMBRA1a C-terminal WD40-like region (reference residues 711-917) in the same proteins. The two regions were aligned independently using MAFFT E-INS-i. Only alignment positions represented by a non-gap, non-ambiguous residue in at least 50% of the sequences are shown. Dark-grey shading marks consensus-matching residues at positions with at least 70% conservation, whereas light-grey shading marks consensus-matching residues at positions with 50-69% conservation. Unshaded residues either differ from the consensus or occur at positions with less than 50% conservation. Each row corresponds to one sequence, ordered as in the phylogenetic tree (Fig. 1a); the coloured bar on the left indicates the taxonomic group (Stramenopiles, orange; Rhodophyta, red; Viridiplantae, green; Metazoa, purple).

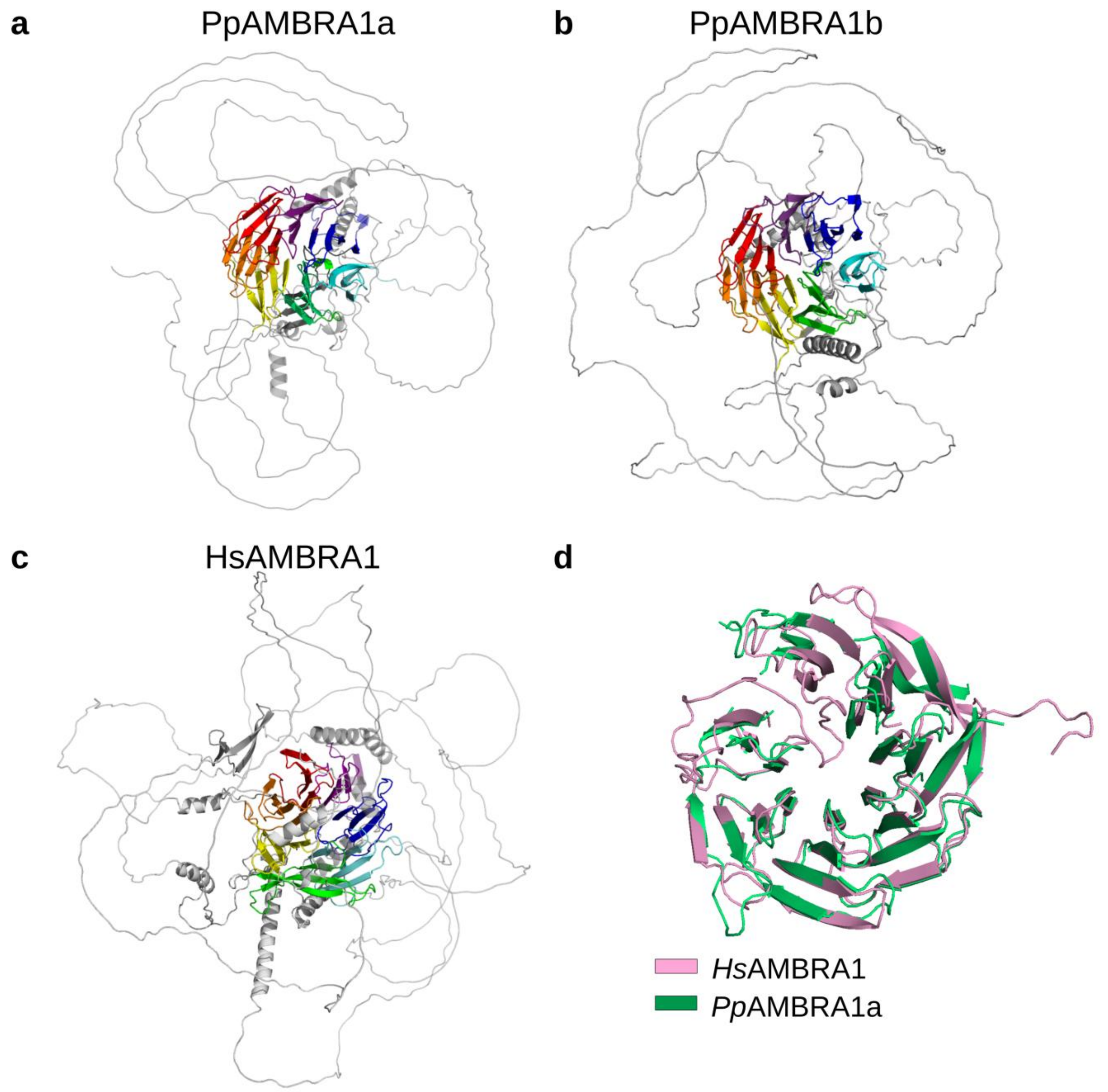


**Supplementary Fig. 2. AlphaFold-predicted structures and structural comparison of *Physcomitrium patens* and *Homo sapiens* AMBRA1.** Predicted structures of (a) *Pp*AMBRA1a (UniProt ID: A0A2K1JYD0, AF-A0A2K1JYD0-F1), (b) *Pp*AMBRA1b (UniProt ID: A0A2K1JGJ8, AF-A0A2K1JGJ8-F1) and (c) *Hs*AMBRA1 (UniProt ID: Q9C0C7, AF-Q9C0C7-4-F1). (d) Structural superimposition of the conserved WD40 core domains of *Pp*AMBRA1a and *Hs*AMBRA1.

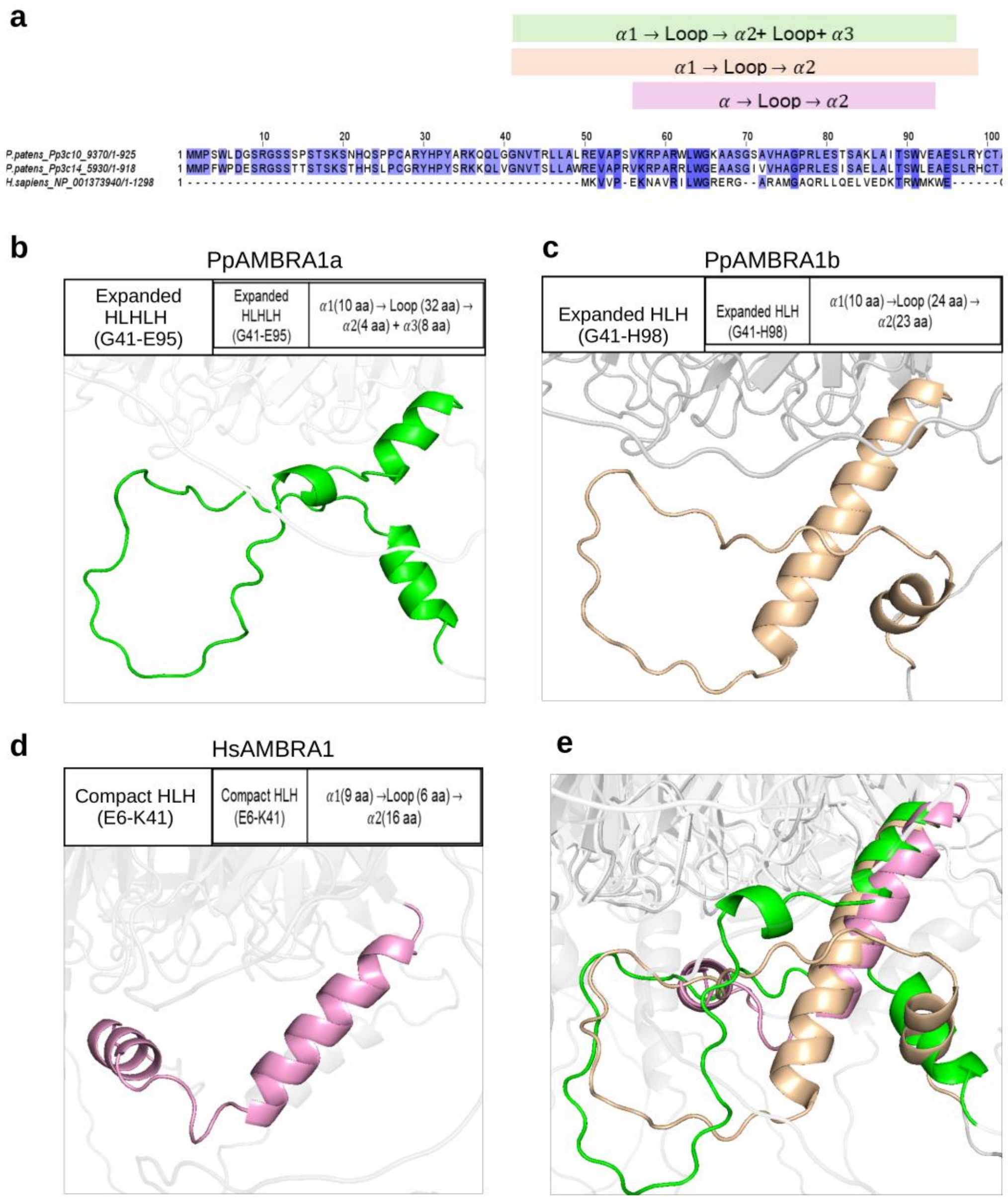


f

| N-terminal structure | RMSD (Å) |
|---|---|
| PpAMBRA1a vs HsAMBRA1 | 0.980 |
| PpAMBRA1b vs HsAMBRA1 | 0.568 |
| PpAMBRA1a vs PpAMBRA1b | 0.229 |

**Supplementary Fig. 3. N-terminal domain of *Pp*AMBRA1A, *Pp*AMBRA1B, *Hs*AMBRA1.** a) Sequence alignment and schematic representation of the secondary structure elements of the N-terminal region from *P. patens* AMBRA1A (green), *P. patens* AMBRA1B (wheat), and *H. sapiens* AMBRA1 (pink). b) Detailed view of the expanded HLHLH motif (G41-E95) of *Pp*AMBRA1A (green), composed of $\alpha1$ (10 aa), a loop (32 aa), $\alpha2$ (4 aa), and $\alpha3$ (8 aa). c) Detailed view of the expanded HLH motif (G41-H98) of *Pp*AMBRA1B (wheat), composed of $\alpha1$ (10 aa), a loop (24 aa), and $\alpha2$ (23 aa). d) Detailed view of the compact HLH motif (E6-K41) of *Hs*AMBRA1 (pink), composed of $\alpha1$ (9 aa), a loop (6 aa), and $\alpha2$ (16 aa). e) Structural superimposition of the N-terminal motifs of *Pp*AMBRA1A, *Pp*AMBRA1B, and *Hs*AMBRA1. f) Root-mean-square deviation (RMSD, in Å) values calculated from the structural alignment of the N-terminal regions across the three proteins.

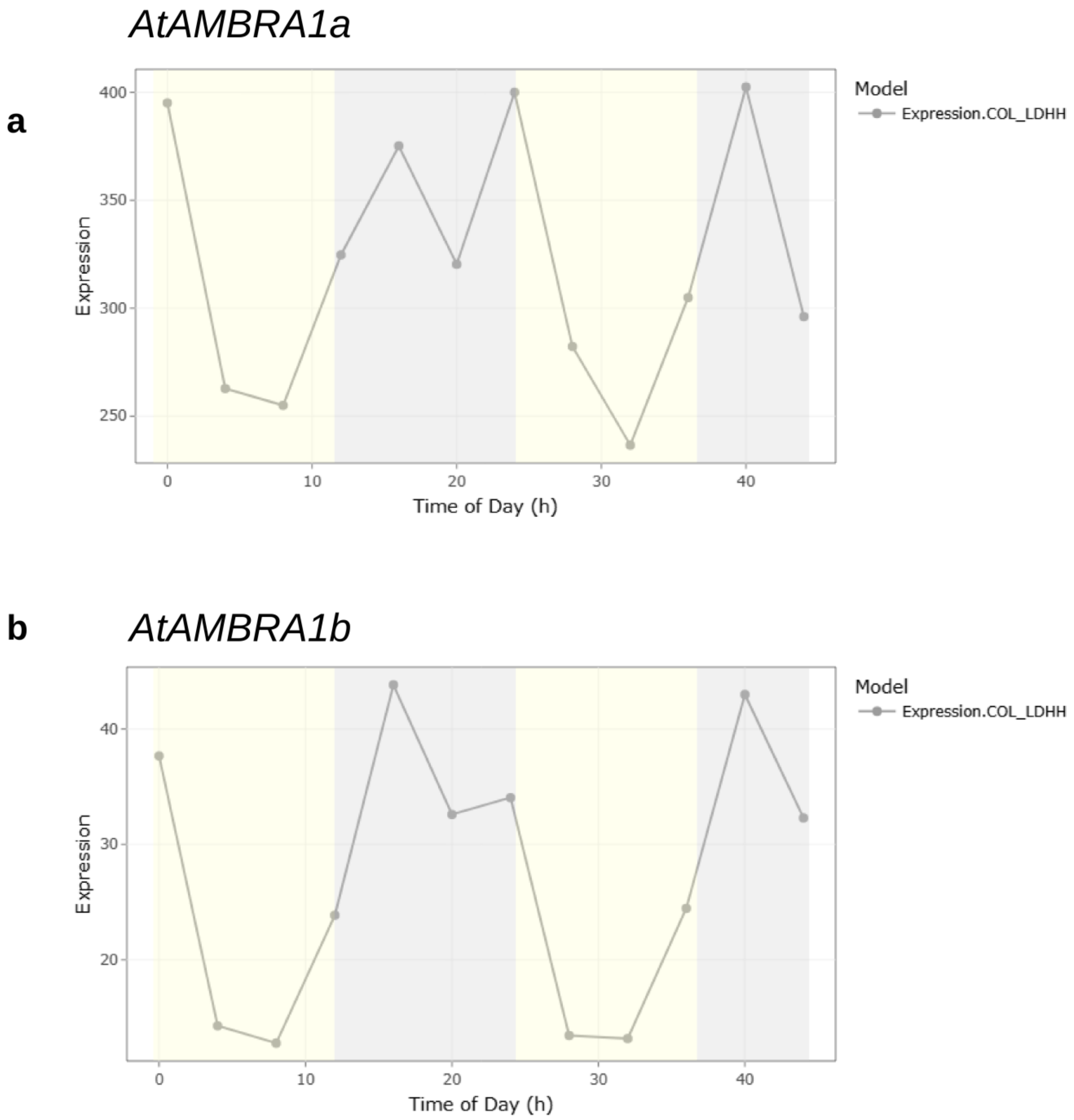


**Supplementary Fig. 4. *Arabidopsis thaliana AMBRA1a* and *AMBRA1b* diurnal expression.** Expression profiles of *AtAMBRA1a* (At1g04140; a) and *AtAMBRA1b* (At5g43930; b) under long-day conditions (COL_LDHH dataset, DIURNAL database). Yellow and grey backgrounds indicate light and dark periods, respectively.

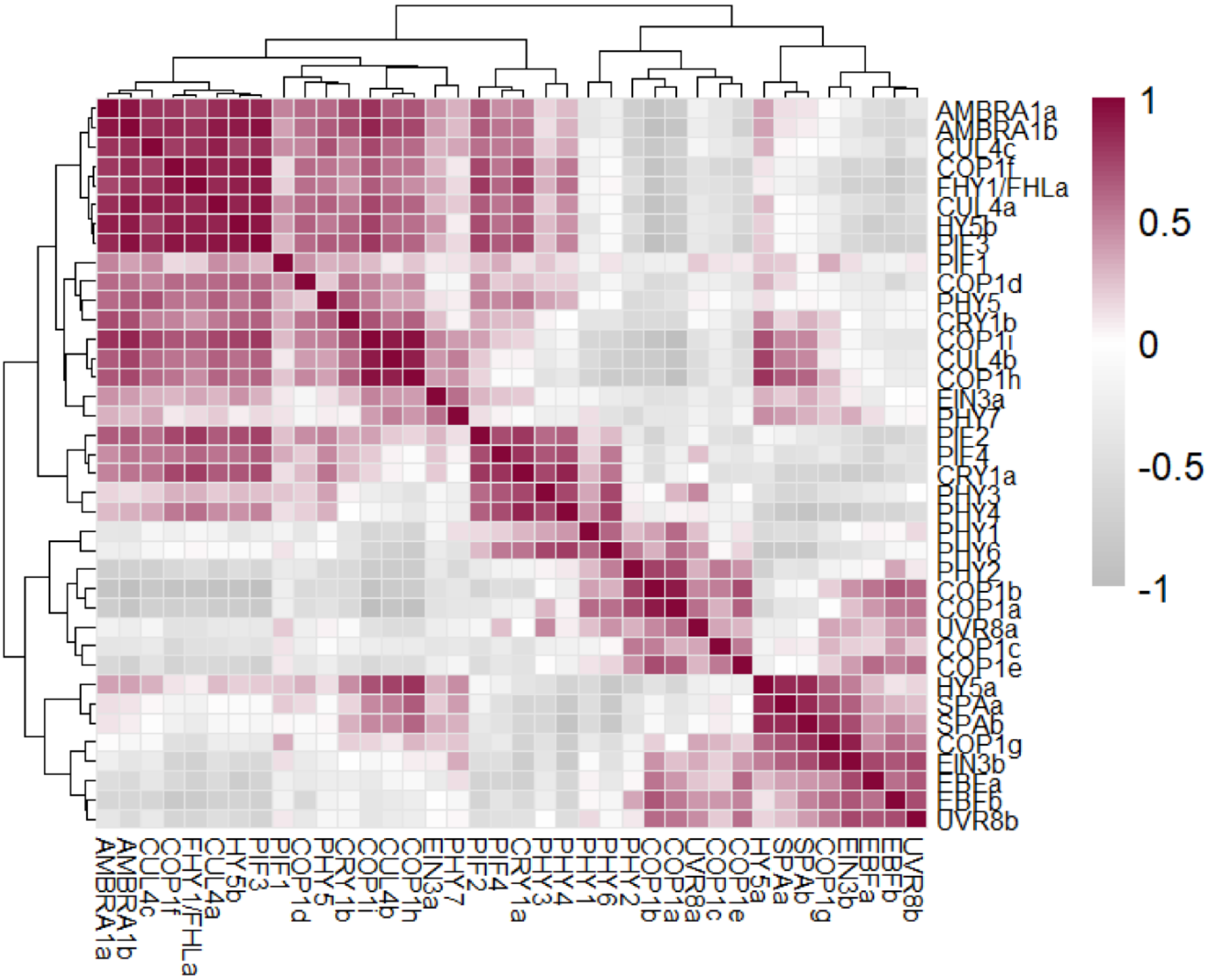


**Supplementary Fig. 5. Correlation matrix between the two *P. patens* AMBRA1 genes and 36 genes defined as part of light signalling core component in [9].** Hierarchical clustering heatmap showing the pairwise Pearson correlation matrix between the two *P. patens AMBRA1* genes (*AMBRA1a* and *AMBRA1b*) and 36 core light signaling genes defined by [9]. The color scale on the right indicates the Pearson correlation coefficient ($r$), ranging from high positive correlation (dark red, $+1$) to inverse correlation (dark gray, $-1$). Both horizontal and vertical dendrograms represent hierarchical clustering based on co-expression profiles.

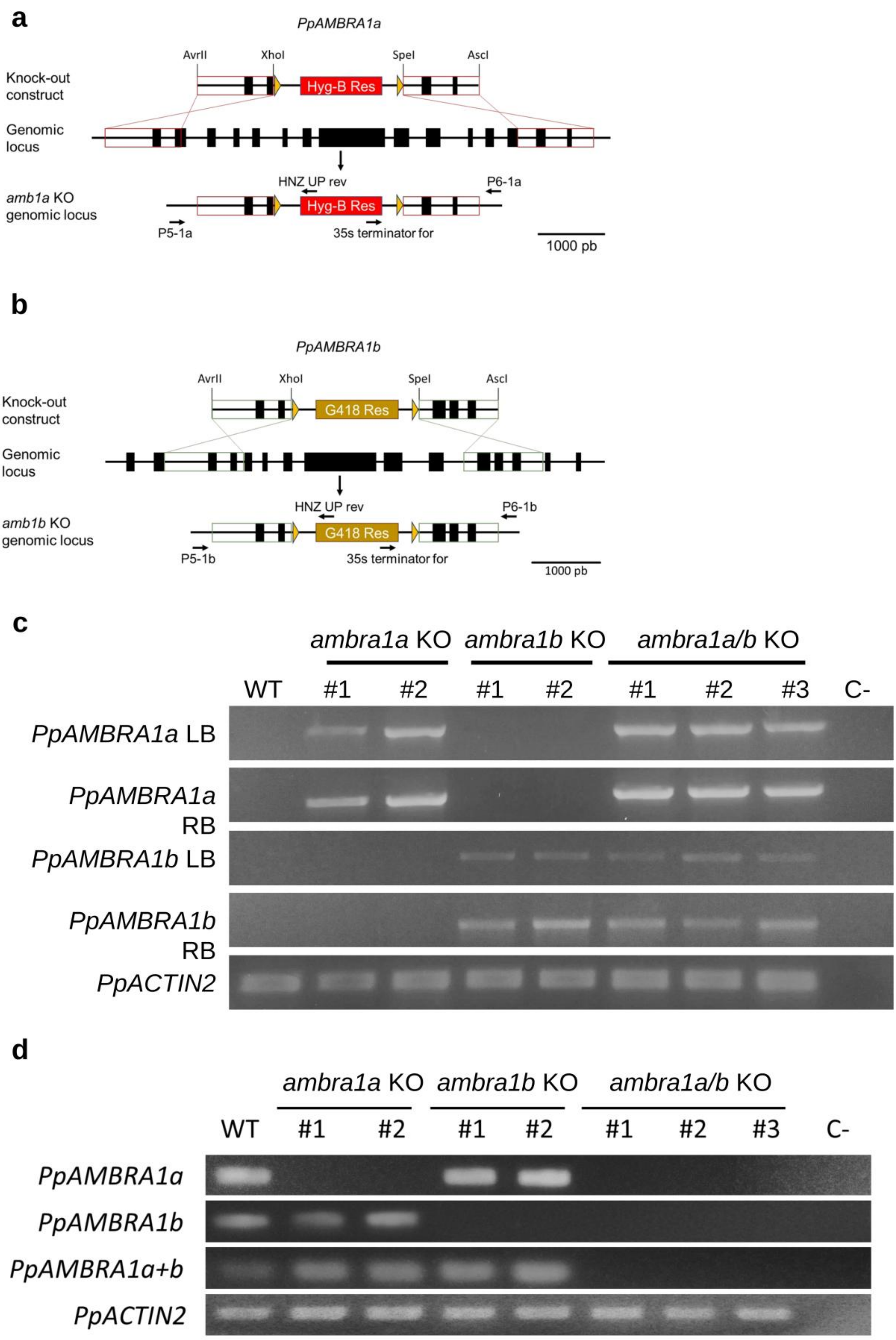


**Supplementary Fig. 6. Generation and molecular validation of *ambra1a* and *ambra1b* deletion mutants.** (a-b) Schematic representation of the strategy used to replace the *AMBRA1A and AMBRA1B* genes with a Hygromycin B resistance cassette (Hyg-B Res) and a G418 resistance cassette (G418 Res). Black boxes represent the exons. The knockout constructs were generated cloning 5' and 3' homology regions (red and green boxes). Restriction enzymes indicated above the constructs represent the sites used for molecular cloning. AvrII and AscI enzymes were also used to linearize the cassette prior to protoplast transformation. Yellow triangles indicate loxP sites flanking the selection marker. Horizontal arrows represent the specific primers used to screen stable resistant lines for site-specific integration of the resistance cassette. (c) Validation of mutant strains by PCR on genomic DNA. Agarose gel electrophoresis confirming the genotype of the wild-type (WT) and transformed lines. LB (Left Border) and RB (Right Border) PCR amplifications demonstrate that the constructs are correctly inserted into the target loci using the primers indicated in (a) and (b). Rows 1-2 show the LB and RB PCRs for the *AMBRA1A* disruption (Hyg-B insertion), while rows 3-4 show the LB and RB PCRs for the *AMBRA1B* disruption (G418 insertion). Row 5 shows the amplification of *ACTIN2* that serves as an internal loading control to confirm uniform DNA template quality across all samples. WT is reported along with two independent *ambra1a* knockout lines, two independent *ambra1b* knockout lines and three independent *ambra1a/b* double knockout lines. C- is the negative control, water was used to substitute genomic DNA for PCR. d) RT-PCR analysis of *PpAMBRA1a* and *PpAMBRA1b* expression in KO lines. To verify the loss of transcript in the generated mutant lines, semi-quantitative RT-PCR was performed on cDNA synthesized from total RNA. Amplification of the *PpAMBRA1a* transcript shows a complete absence of expression in *ambra1a* KO and *ambra1a/b* KO lines, while expression remains unaffected in the WT and ambra1b KO lines. Amplification of the PpAMBRA1b transcript confirms its absence in ambra1b KO and *ambra1a/b* KO lines, whereas it is normally expressed in WT and *ambra1a* KO backgrounds. Use of a primer pair targeting a conserved region of both paralogs (PpAMBRA1a+b) results in amplicons in the single KO lines (representing the remaining active paralog) but shows a total absence of signal in the three independent *ambra1a/b* double KO. *PpACTIN2* was used as an internal constitutive control to ensure equal cDNA loading across all samples. C- indicates the negative control to exclude DNA contamination in the PCR experiment.

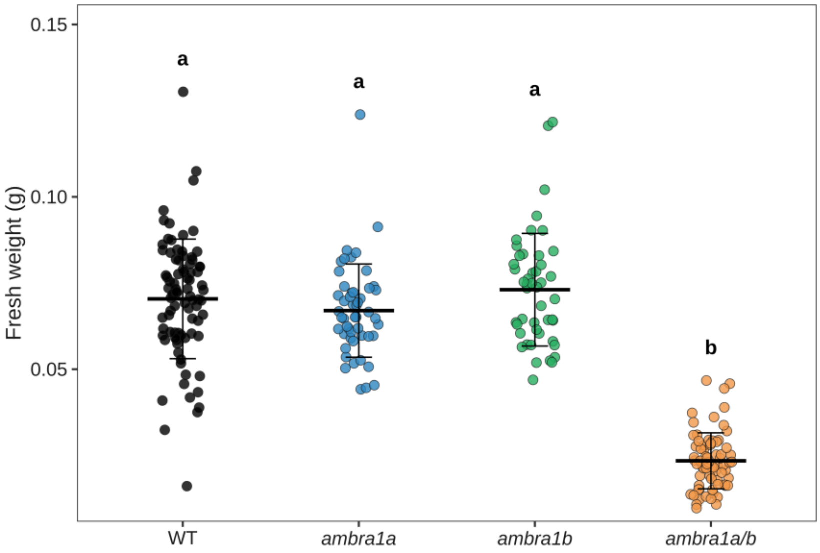


**Supplementary Fig. 7. Phenotypic characterization of *ambra1a/b* lines under long day conditions.** Fresh weight of WT, *ambra1a*, *ambra1b* and *ambra1a/b* colonies grown for 28 days under standard photoperiod.

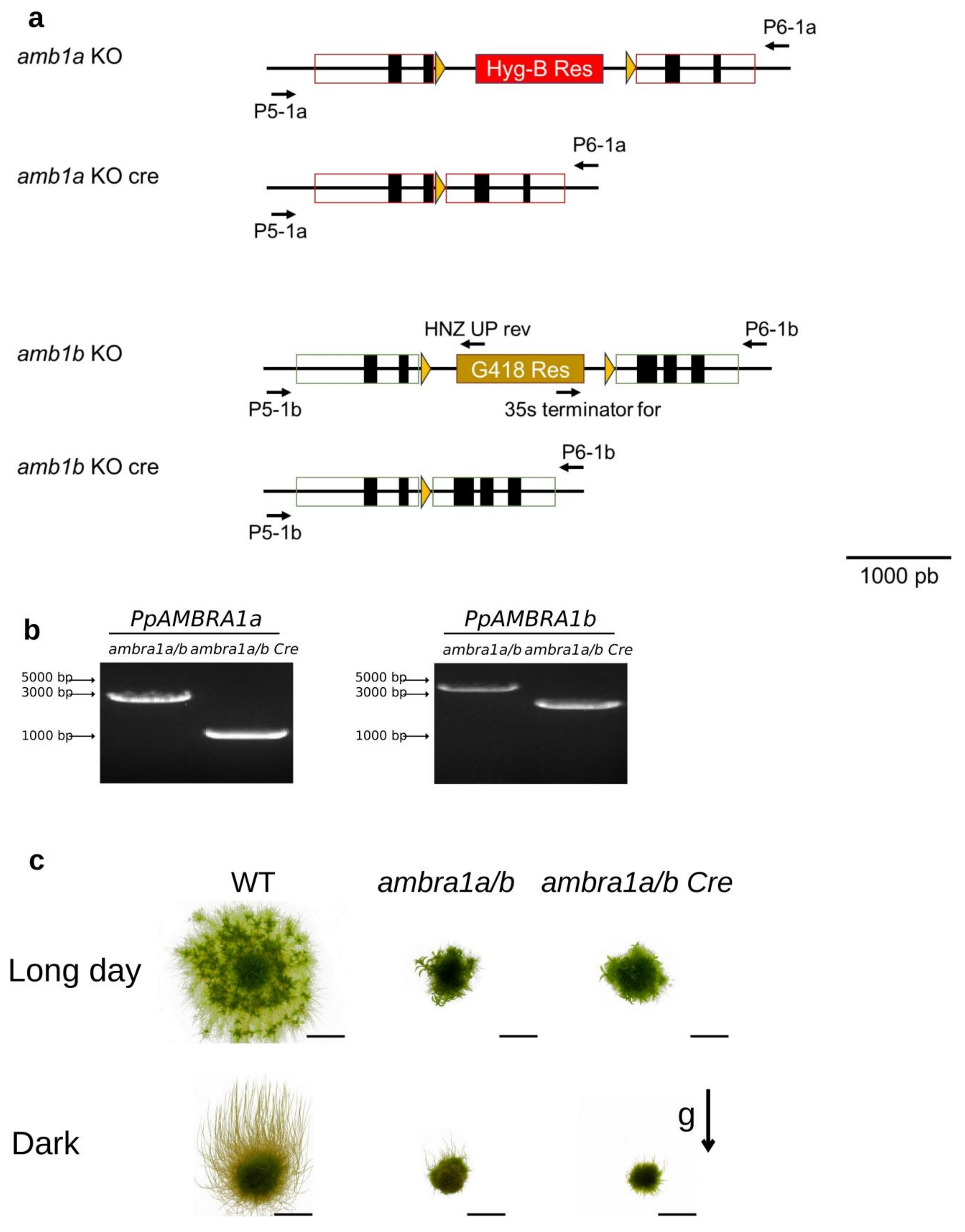


**Supplementary Fig. 8. Cre-mediated excision of selection markers in AMBRA1 mutant lines.** a) Schematic representation of the genomic loci before and after Cre-recombinase treatment. The selection markers, Hyg-B Res (red) and G418 Res (orange), were originally integrated to disrupt the AMBRA1a and AMBRA1b genes (exons indicated by black boxes). These cassettes are flanked by loxP sites (yellow triangles). Following transient expression of Cre recombinase, site-specific recombination between the loxP sites results in the excision of the resistance genes, leaving behind a single loxP scar. The cloned homology regions are indicated by red (AMBRA1a) and green (AMBRA1b) open boxes. Horizontal arrows indicate the position of the primers used for PCR screening. Scale bar represents 1000 bp. b) PCR verification of marker excision. Agarose gel electrophoresis shows the shift in band size following the successful removal of the resistance cassettes. In the *ambra1a/b* double KO lines before Cre treatment, the amplicons for AMBRA1a and AMBRA1b loci are larger due to the presence of the resistance markers (~2.5-3 kb). After Cre-mediated excision (*ambra1a/b* Cre), the PCR products show a significant reduction in size (to ~0.5-1.2 kb), corresponding to the expected "clean" deletion of the target regions. A DNA ladder is provided on the left with sizes indicated in base pairs (bp). c) Growth phenotype of *ambra1a/b* Cre under standard photoperiod and continuous darkness.

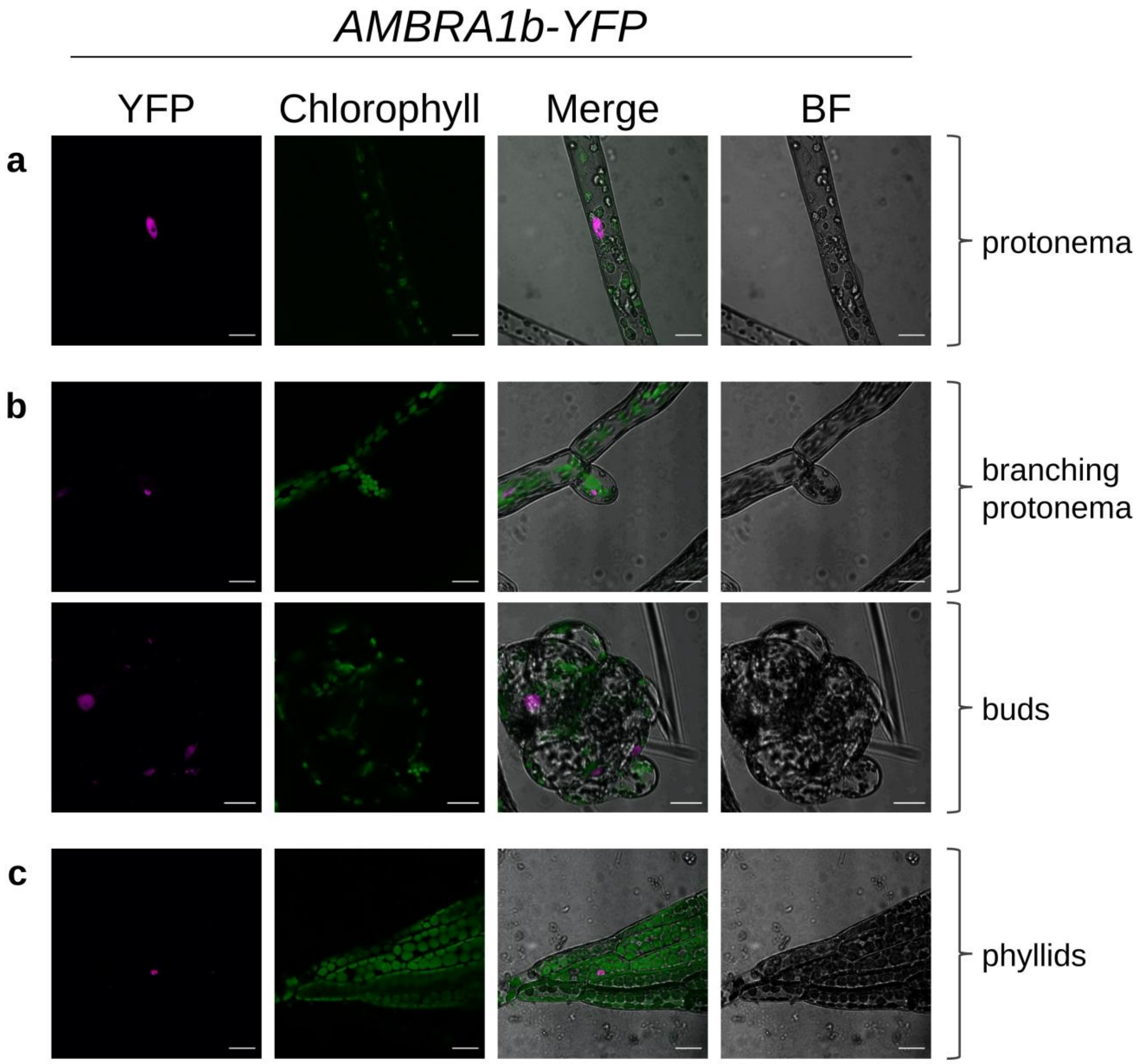


**Supplementary Fig. 9. Subcellular localization of AMBRA1b-YFP in (a) protonema, (b) branching protonema and buds, and (c) phyllids.** Scale bar= 5µm.

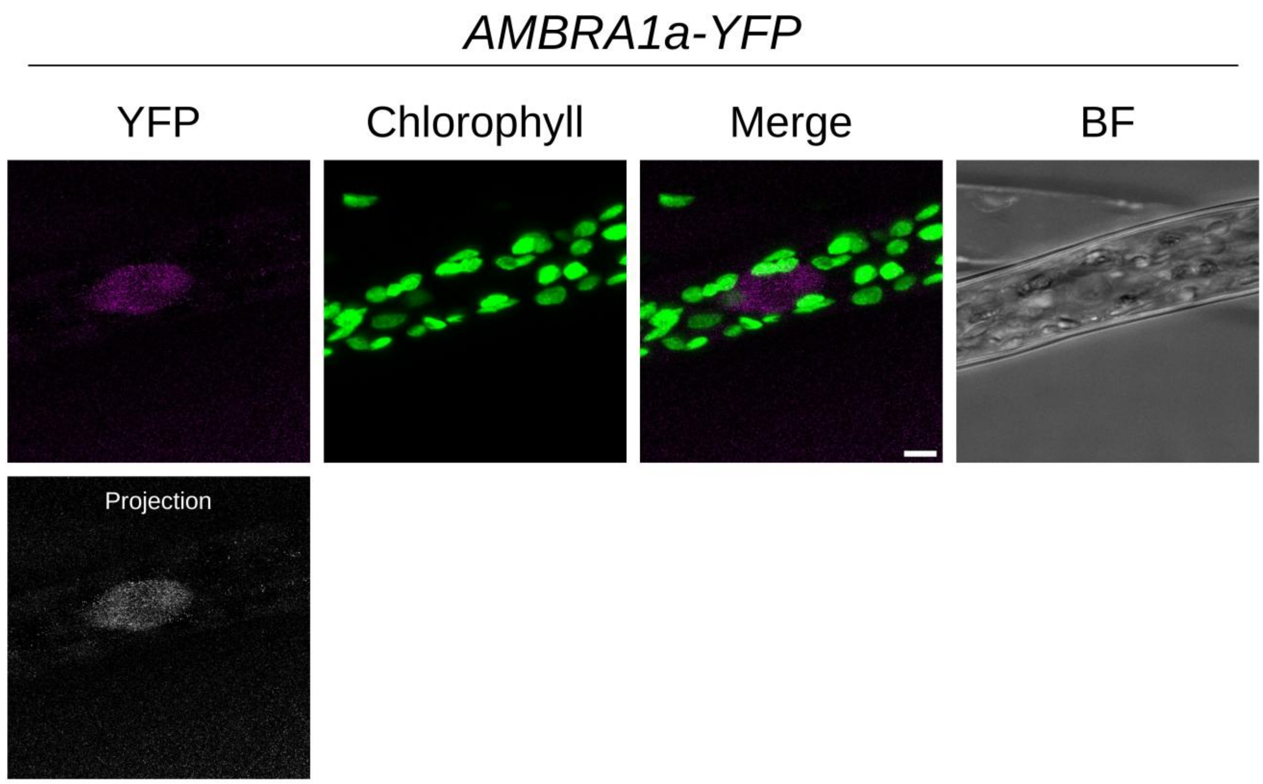


**Supplementary Fig. 10. Subcellular localization of AMBRA1a-YFP in protonemal cells.** YFP signal (magenta), chlorophyll autofluorescence (green), merged and brightfield (BF) images are shown; the lower panel shows a maximum-intensity projection of the YFP signal. Scale bar = 5 µm.

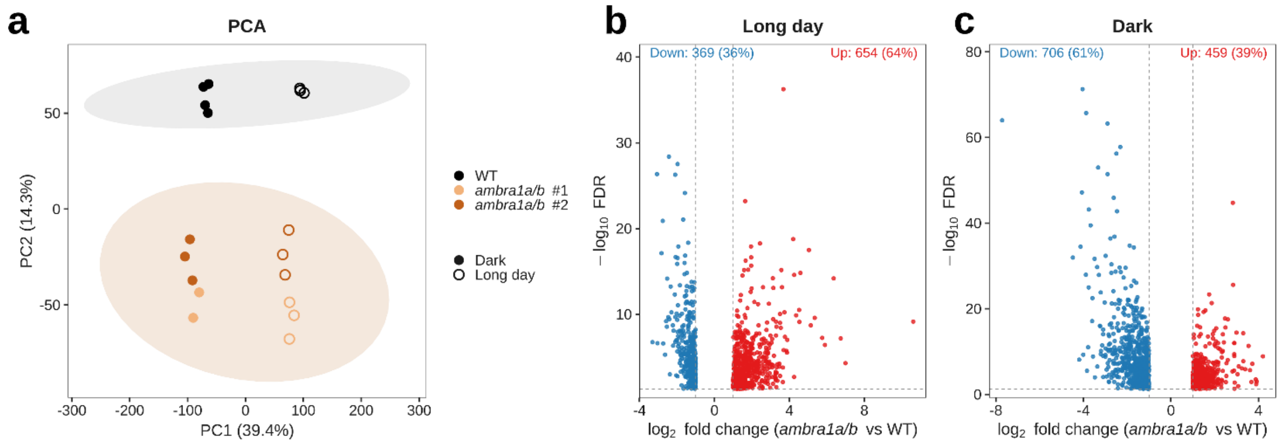


**Supplementary Fig. 11. Transcriptome profile of WT and *ambra1a/b* KO plants grown under a long-day photoperiod or in continuous darkness.** a) Principal component analysis (PCA) of WT and *ambra1a/b* KO (clone 1 and clone 2) transcriptomes under long-day and dark conditions; dashed ellipses indicate the 95% confidence interval for each genotype. b, c) Volcano plots of differentially expressed genes (DEGs) in *ambra1a/b* vs WT under long-day (b) and dark (c) conditions (|log2FC| > 1, FDR < 0.05); percentages of up- and downregulated genes are indicated above each plot.

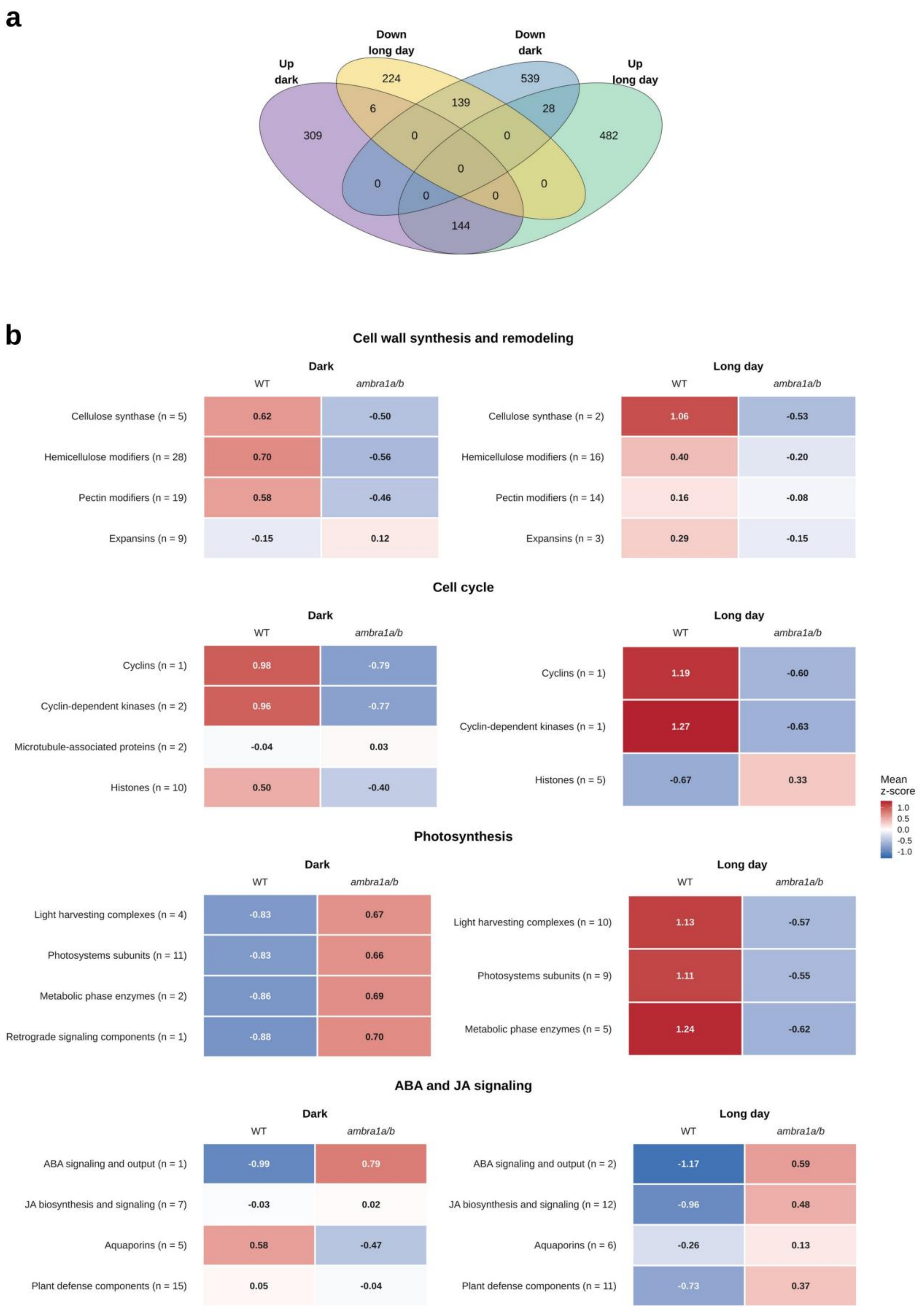


**Supplementary Fig. 12. Transcriptional response of core DEGs in *ambra1a/b* mutants under long-day and dark conditions.** a) Four-way Venn diagram of the differentially expressed genes (DEGs, *ambra1a/b* vs WT, FDR-filtered) identified in dark- and light-grown plants, split by direction of change: UpDark (n = 459), DownDark (n = 706), UpLight (n = 654), DownLight (n = 369). b) Mean expression z-scores of DEGs grouped into four functional categories: cell wall synthesis and remodeling, cell cycle, photosynthesis, and ABA/JA signaling. Genes were assigned to each category and subcategory (rows; n = number of member genes detected among the DEGs) based on gene symbol and functional annotation (Phytozome v3.3 annotation / Arabidopsis best-hit description). For each gene, log2(CPM+1) values were row-scaled (z-score) across all samples and then averaged within genotype (WT, *ambra1a/b*) and light regime (Dark, Long day). Color scale: mean z-score, with blue indicating expression below the gene's own across-sample mean and red indicating expression above it.

**List of Supplementary Tables and Movie**

Supplementary Table 1. Sequences used for phylogenetic tree reconstruction.

Supplementary Table 2. Co-expression correlation matrix of *PpAMBRA1* and photomorphogenic genes.

Supplementary Table 3. DeepLoc subcellular localization predictions.

Supplementary Table 4. Differentially expressed genes (DEGs) in *ambra1* mutants under Long-Day (LD) conditions.

Supplementary Table 5. Differentially expressed genes (DEGs) in *ambra1* mutants under Dark (D) conditions.

Supplementary Table 6. GO and KEGG pathway enrichment analyses under Long-Day (LD) and Dark (D) conditions.

Supplementary Table 7. Shared DEGs between Long-Day and Dark conditions.

Supplementary Table 8. Genes with opposite regulation between Long-Day and Dark conditions.

Supplementary Table 9. Shared DEGs between *ambra1* and *cop1_9x* mutants under Long-Day (LD) conditions.

Supplementary Table 10. Shared DEGs between *ambra1* and *cop1_9x* mutants under Dark (D) conditions.

Supplementary Table 11. GO and KEGG enrichment analyses of concordantly and discordantly regulated genes in *ambra1* and *cop1_9x* mutants.

Supplementary Table 12. List of oligonucleotide primers used in this study.

Supplementary Movie 1. Nuclear YFP signal distribution during cell division into two daughter cells.
